# Multiscale mechanics and temporal evolution of vimentin intermediate filament networks

**DOI:** 10.1101/2021.01.30.428887

**Authors:** Anna V. Schepers, Charlotta Lorenz, Peter Nietmann, Andreas Janshoff, Stefan Klumpp, Sarah Köster

## Abstract

The cytoskeleton, an intricate network of protein filaments, motor proteins, and crosslinkers, largely determines the mechanical properties of cells. Among the three filamentous components, F-actin, microtubules, and intermediate filaments (IFs), the IF network is by far the most extensible and resilient to stress. We present a multiscale approach to disentangle the three main contributions to vimentin IF network mechanics – single filament mechanics, filament length, and interactions between filaments – including their temporal evolution. Combining particle tracking, quadruple optical trapping and computational modeling, we derive quantitative information on the strength and kinetics of filament interactions. Specifically, we find that hydrophobic contributions to network mechanics enter mostly via filament elongation kinetics, whereas electrostatics have a direct influence on filament–filament interactions. These results indicate that cells might need to explicitly suppress attractive interactions to re-organize the extremely stable cellular vimentin network.

## Introduction

Cells have remarkably diverse and dynamic mechanical properties that are largely determined by the cytoskeleton [1]. The adaptability of the cytoskeleton relies on the unique and distinct mechanical properties of the different biopolymers that form this composite network [2]. Intermediate filaments (IFs) are particularly interesting from a mechanical point of view. By contrast to F-actin and microtubules, both single IFs and networks thereof display an enormous extensibility and stability at high strains. These properties have been impressively shown for vimentin IFs, which are found in cells of mesenchymal origin [2–4]. Rheological studies reveal that vimentin networks are soft at small strains, but show pronounced strain stiffening up to a critical strain, at which the networks soften [2, 5–8], unless they are covalently cross-linked [8]. Furthermore, vimentin networks stiffen in the presence of divalent ions, which has been attributed to transient interactions between filaments [6–10]. Whereas these previous studies provide a rich pool of information on network mechanics, they cannot unravel the effects of filament elongation kinetics, single filament mechanics and filament–filament interactions, which are all superimposed. For instance, an influence of divalent cations for IF network mechanics is indisputable, but the exact mechanism by which they alter the network properties remains elusive.

Here, we present a targeted experimental and computational approach that enables us to disentangle the contributions of single filament properties and filament–filament interactions to network mechanics. With our multiscale approach we extract binding and unbinding rates for single interactions and thereby directly quantify the transient nature of direct interactions between vimentin filaments.

## 1 Results and Discussion

### 1.1 Vimentin filament networks mature and stiffen on time scales of days

Previous studies of vimentin network mechanics have typically focused on time points of about 1 - 2 h after initiation of filament assembly and network formation, which in most experimental protocols occur simultaneously [5–7, 9, 10]. We extend this work and study the formation of vimentin IF networks and the evolution of the network mechanics over a broad time range, from 3 h to 144 h after initiation of assembly, and at standard conditions, *i.e.* in the presence of 100 mM KCl. We record tracks of microparticles embedded in the networks, as shown in Fig. 1a (center), and calculate the mean squared displacement (MSD) for each assembly duration, see Fig. 1b (center). At 3h (black line), microparticle tracking (MPT) reveals subdiffusive behavior. For comparison, the MSD for diffusive motion is shown as the purple dashed line. For longer network formation times, we observe a stiffening of the vimentin network (for color code see legend). The decreased mobility of the particles leads to a flattening of the MSDs at long lag times; as expected, after very long network formation times, the beads are confined and the MSDs approach a plateau.

**Figure 1.**
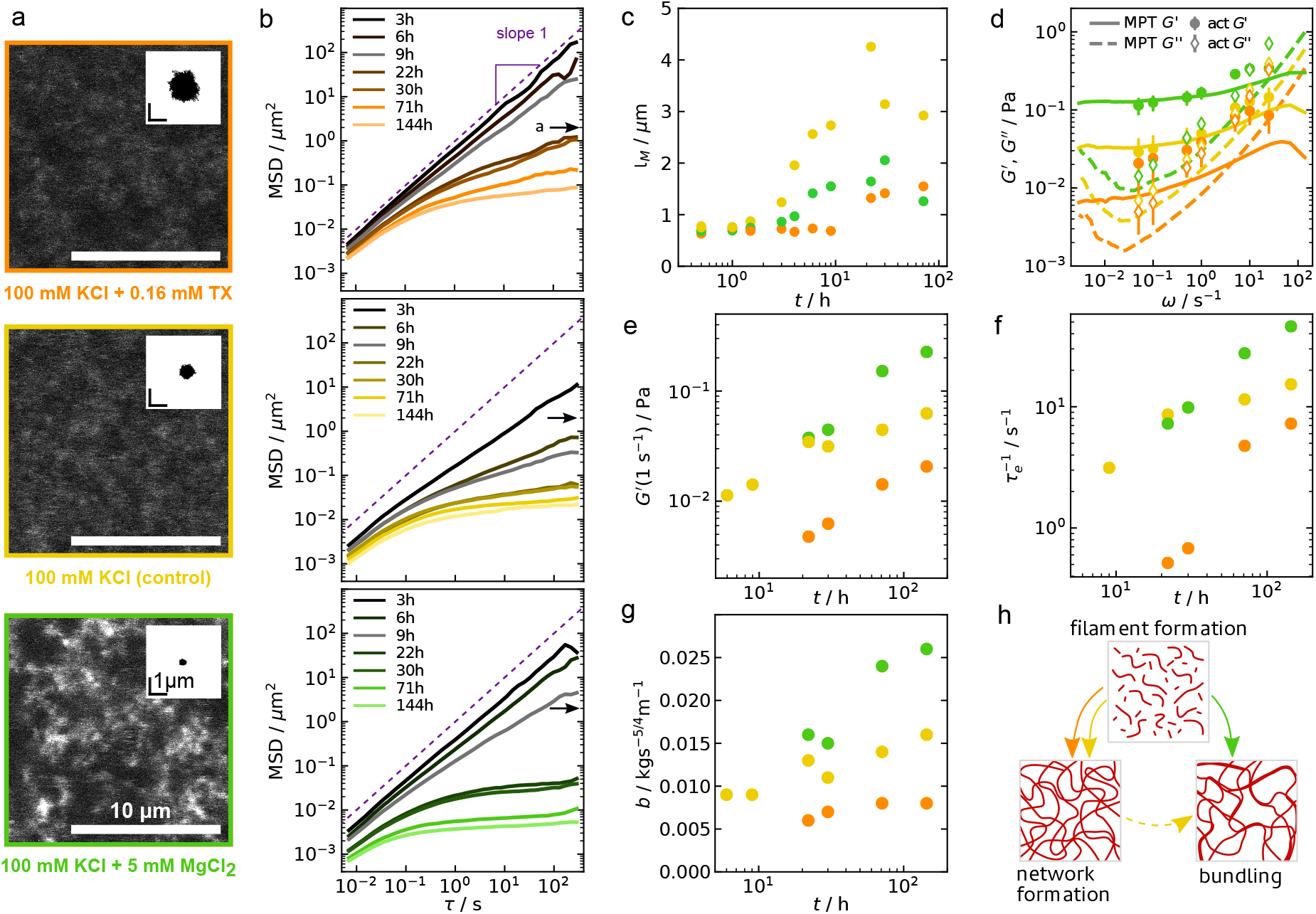
Formation and mechanics of vimentin networks. a) Microscopy images of vimentin networks after 3 days (74 h) of network formation time, buffer conditions indicated below each image. The insets show representative traces from MPT, recorded over the course of 6 min after three days (71 h) of network formation. b) Median MSD for each condition for increasing network formation times (for color code, see legend), the particle diameter *a* = 2*μ*m is indicated by arrows. Individual curves are shown in Figs. S5-S7. c) Elongation of filaments. The length average *I_M_* is calculated from step growth polymerization modeling of filament length histograms shown in Fig. S3. d) Comparison of active (symbols) and passive (lines) microrheology measurements after 3 days. The error bars denote the standard deviation of the active measurements. e) Temporal evolution of *G′* at *ω* = 1 s^-1^. The values for the median curves are shown. f) Temporal evolution of the relaxation times of single filaments in the network. g) Temporal evolution of the bundling parameters, calculated from the shift of *G″ ≈ bω^3/4^* at high frequencies. h) Schematic of the filament and network formation process with the competition between lateral assembly, i.e. bundling, and filament elongation that leads to an entangled network.

The stiffening of the network continues over the whole assembly duration probed. From 6h on, the particles are confined, as the movement does not exceed the particle diameter of *a* = 2 *μ*m after the total measuring time. This transition from subdiffusive to confined motion is caused by the slowly growing filaments. The hierarchical assembly pathway of vimentin filaments and the corresponding time scales are shown in Fig. S1. To characterize the network further, we determine the minimum length for entanglement by calculating the length average of the filaments, *l_M_*, according to Flory’s theory [11], see Fig. 1c and Figs. S2-S4, which reaches ≳1.2*μ*m between 3-6h.

#### The filament length depends on elongation and lateral association

To investigate the cause of the network stiffening, we analyze the temporal evolution of the filament length. Specifically, the formation kinetics, architecture and mechanics of the networks depend on a careful balance between the slow longitudinal assembly of single filaments and the lateral association of filaments, *i.e.* bundling. This balance is influenced by lateral interactions between the filaments [12] which we examine by varying the interaction strength, specifically hydrophobic and electrostatic interactions between vimentin IFs [5, 13, 14]. For keratin IFs, in particular, it has been observed that the suppression of hydrophobic interactions by detergents may cause softening of IF networks [13, 15]. By contrast, an increase of electrostatic attractions, mediated by divalent ions, leads to a stiffening of IF networks [6, 8–10]. To separate hydrophobic and electrostatic effects, we assemble vimentin filaments in the presence of additional non-ionic detergent TritonX-100 (TX) at 0.16 mM to diminish hydrophobic interactions, or 5mM MgCl_2_ for increased electrostatic attraction, see Fig. 1c. We observe a slower filament elongation with overall shorter filaments in both cases. In the presence of TX, suppression of hydrophobic interactions severely impairs the elongation (Fig. S3). We therefore conclude that hydrophobic interactions within filaments, *i.e.* between subunits, play a pivotal role during vimentin IF assembly. For added MgCl_2_, we follow the line of arguments of Ref. [12] and assume that the balance is shifted towards increased attraction between filaments, leading to lateral association of short filaments rather than longitudinal annealing. For vimentin IFs, such bundling events have been directly observed in the presence of sufficiently high concentrations of MgCl_2_ [16, 17].

### 1.2 Electrostatic and hydrophobic interactions lead to mechanically distinct networks

The bundle formation caused by the addition of MgCl_2_ not only influences elongation kinetics, but also network mechanics. We extend previous studies of vimentin network mechanics in the presence of divalent ions [6, 8–10] to much longer assembly time scales, since we expect a saturation of the network mechanical properties after days rather than hours, as shown above for the control condition. We assemble the networks in the same buffer conditions as the single filaments, with 0.16 mM TX for reduced hydrophobic interactions, Fig. 1a (top), or 5mM MgCl_2_ for increased electrostatic attraction (bottom). Confocal images of the respective networks reveal no apparent difference between the control condition and in the presence of TX, whereas the networks are more heterogeneous in the presence of MgCl_2_.

As expected from the filament elongation studies, the corresponding MPT data in Fig. 1b, top and bottom, show that the network formation, in particular the transition from sub-diffusive to confined motion, is delayed for both conditions compared to the control. As we eventually observe confinement, we reason that the relevant length scales of the networks are comparable. We transfer the MSDs that show confinement into the complex shear modulus *G\** = *G′* + *iG″* with the storage *G′* and loss *G″* moduli, as shown in Figs. S8-S10 [18]. Analysis of the plateau modulus *G′*(1s^-1^) illustrates the continuous stiffening that all samples undergo with advancing time, see Fig. 1e. From 22 h on, the network with MgCl_2_ is the stiffest, the network in the control sample shows intermediate stiffness and the sample with detergent stays the softest throughout. As an example, the comparison of *G′* and *G″* of the networks after 71 h in Fig. 1d illustrates this difference clearly. Active microrheology by optical trapping of microparticles embedded in the respective samples (Fig. 1d and Fig. S11), validates the passive microrheology results from MPT with remarkable agreement.

The connectivity of the networks influences the frequency dependence of the plateau. In presence of MgCl_2_, see Fig. S12, the plateau is the least frequency dependent, indicating a more pronounced contribution of cross-linking rather than entanglement to the network mechanics [10]. This is in agreement with the established hypothesis that divalent ions serve as transient cross-linkers in IF networks [6, 8, 9]. The binding and unbinding rates of these transient cross-links can, however, not be inferred from our measurements. The increased connectivity of the networks as well as stiffening due to bundle formation further causes an increase of the inverse relaxation time 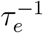, as determined from the cross-over frequency of *G′* and *G″*, see Fig. 1f [19]. Above 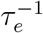 single filament or bundle mechanics dominate over the network properties (see Fig. S13).

As we expect an influence of bundling on the network mechanics, we analyze the coupling of filaments in bundles by investigating the slope of G* at high frequencies. At *ω >* 5 s^-1^, G″ follows a power law with exponent 3/4 for all samples, see Eq. (5) and Fig. S14, as theoretically predicted for networks consisting of single filaments or bundles that are coupled so strongly that they behave like single filaments [20].

Furthermore, the offset of the power law regime of *G″*, *b*, see Eq. (6), indicates trends in the degree of bundling, as it strongly depends on the persistence length of the filaments *l_P_* or bundles *l_P(bundle)_ = N^2^l_P(filament)_*. Fig. 1g shows a strong increase of b over the network formation times in the presence of Mg^2+^, and a much weaker increase for the control condition or in the presence of TX, suggesting that in the presence of Mg^2+^ there are, on average, more filaments per bundle, see also Fig. S14. All processes described above are shown schematically in Fig. 1h.

#### Single filament mechanics are unaffected by detergents or divalent ions

Apart from filament elongation kinetics, single filament mechanics and filament–filament interactions may be influenced by the addition of cations or detergent. Indeed, MPT probes all of these effects in a combined fashion. To disentangle these effects, we focus on single filament mechanics next. The mechanical properties of single vimentin IFs are well established [4, 21–23] and it has been shown that they sensitively depend on the surrounding charges [24]. We find that divalent ions do not stiffen single filaments and TX slightly softens filaments, as shown in Fig. S15. We can therefore safely exclude single filament mechanics as the main cause for the marked differences found for the networks in the three buffer conditions.

### 1.3 Electrostatics increase single filament–filament interactions

After investigating filament elongation and single filaments mechanics, in a third step, we scrutinize filament–filament interactions with single molecule precision. To this end, we bring two filaments in contact in a perpendicular configuration using a quadruple optical trap set-up, see Fig. S16. Confocal images of the experiment are shown in Fig. 2a. We slide one filament perpendicularly along the other one and measure the interaction force by the deflection of the particle “b1”, marked green in Fig. 2b. The resulting interaction force for the filament pair shown in Fig. 2a is plotted in Fig. 2c. We extract the duration of the interaction (red in Fig. 2c) and the strength of the interaction from the maximum interaction force, *F_i_*, before the interaction ruptures, see also Fig. S17. Notably, not all filament pairs show this interaction behavior, and some pairs show no detectable interaction at all. In other cases, the filaments interact, the interaction ruptures and the filaments start to interact again before the force has dropped to zero, see Fig. 2d. Lastly, filaments can interact strongly, see Fig. 2e and Fig. S18, which, only in the presence of MgCl_2_, leads to remarkable “zipping events”, *i.e.* the parallel association of two filaments [25] as shown in Fig. 2f and Fig. S19. From these events, we conclude that interactions are localized and do not slip.

**Figure 2.**
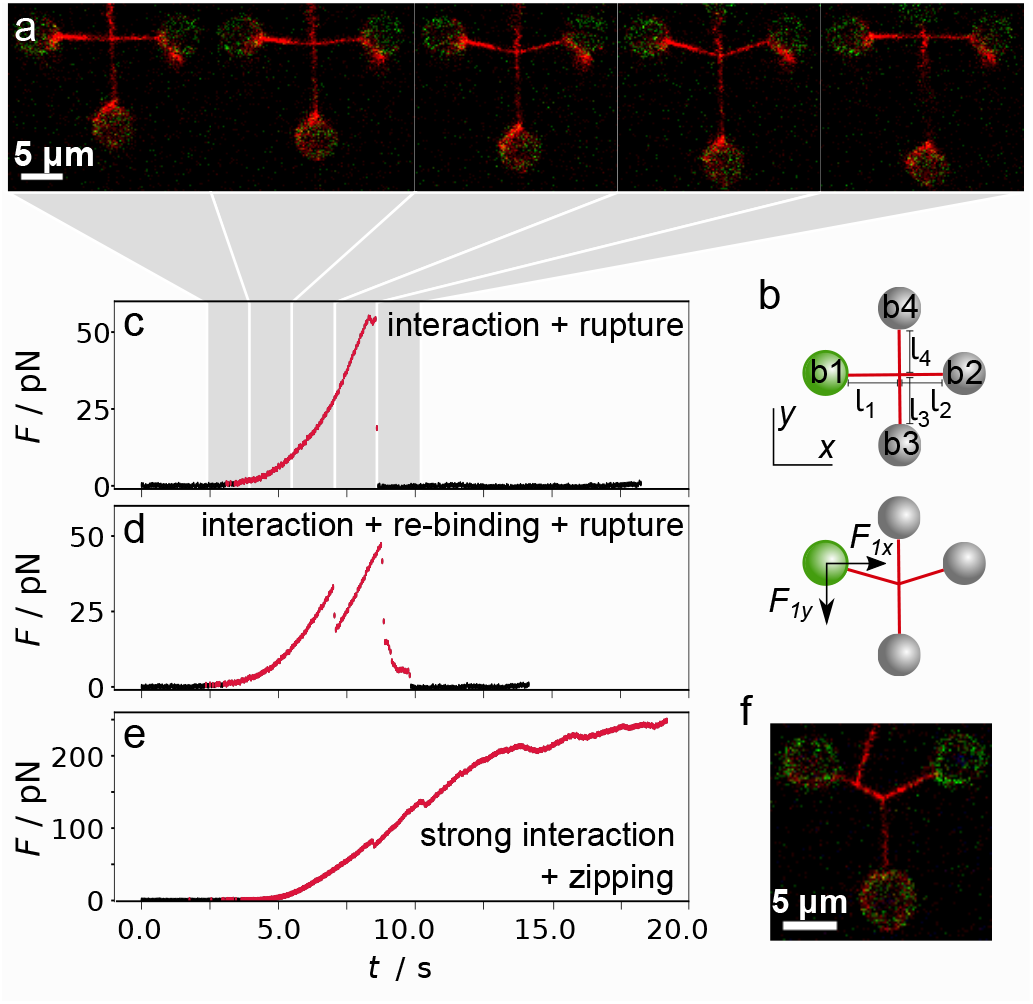
Interaction measurements. a) Confocal image series of a filament pair showing deformation of the horizontal filament as a consequence of the interaction and the return to the initial conformation after rupture of the interaction. b) Schematic of the measuring geometry and nomenclature used in the experiments. We measure and analyze the force *F_1y_* acting on b1 (green). c) Typical interaction force, *F*, plotted against time for the filament pair shown in a); rupture force *F_i_*. d) Typical force-time curve for a measurement with interaction, rupture of the interaction with rebinding under force (kink) and rupture of the interaction. e) Force-time curve for a strong interaction of filaments that show ‘zippinG′, as shown in the micrograph in f).

As we perform the optical trapping experiments in a microfluidic flow cell, we can readily exchange the measuring buffer. A schematic of the experiment is shown in Fig. S16. For the interaction measurements, we always assemble filaments in standard assembly buffer and then move both filaments into measuring buffer, where we bring the filaments in contact and record a sliding assay. This allows us to measure the interactions in extreme buffer conditions, such as 20mM MgCl_2_ where filament networks collapse [16, 17], or high detergent concentrations *c*(TX) = 1.6 mM.

The distribution of forces *F_i_* shown in Fig. 3a at both TX concentrations (red, 1.6 mM and orange, 0.16mM) are similar to the control condition (yellow). We calculate the force-independent binding rate *r_e,b_* as the ratio of the number of interactions that start at zero force and exceed 1 pN, and the experiment time during which we observe no interaction. Since TX, compared to the control condition, has no effect on two parameters that largely determine network stiffness, the strength of the interactions and *r_e,b_*, see Fig. 3b, we conclude that the relative softness of vimentin networks in presence of TX is predominantly caused by impaired filament elongation. It is, however, remarkable that the filaments do interact with non-negligible forces of up to 57pN (1.6mM TX), 85 pN (0.16 mM TX) and 112pN (control). Contrary to observations for keratin networks [13, 15], suppression of hydrophobic interactions does not affect the attraction between single vimentin filaments. Therefore, even without divalent ions present, the filaments interact with physiologically relevant forces [26–28].

**Figure 3.**
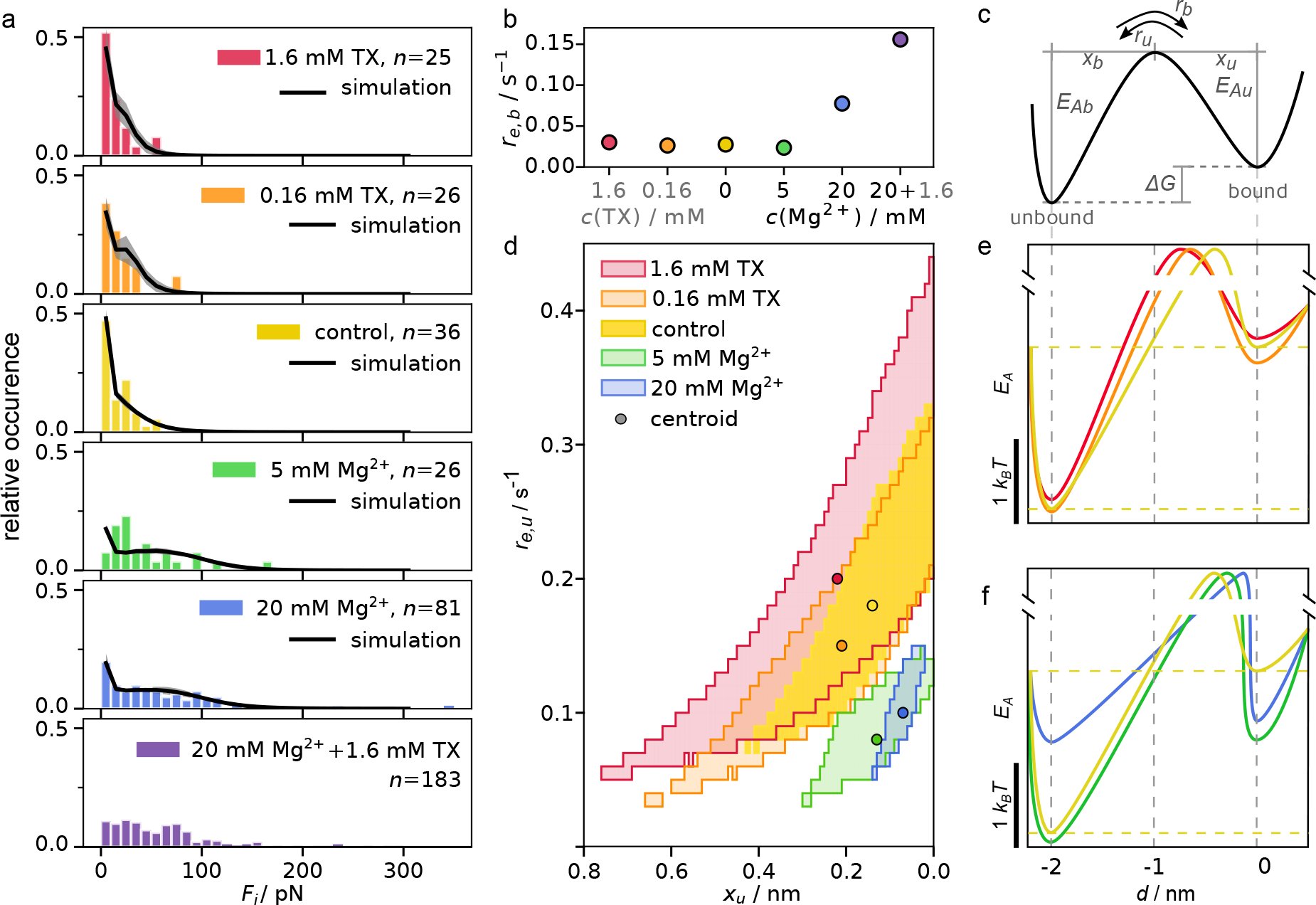
Results of the interaction experiments and simulation. a) Histograms of the rupture forces from experiments (colored bars) and mean curve of the simulation results (black) with standard deviation (shaded area). With decreasing TX concentration (red, orange, yellow), higher forces are reached. The addition of Mg^2+^ (green, blue, purple) causes a broadening of the force distribution and a shift towards higher forces. b) Force-independent binding rate *r_e,b_*. The binding rate is constant, unless Mg^2+^ is present at a concentration of 20 mM. c) Schematic of the two state model used for the simulations. d) Parameter pairs of *x_u_* and the force-independent unbinding rate *r_e,u_*, extracted from the simulation, that satisfy the 5 % significance level in the Kolmogorv-Smirnov test. The centroid is marked. e),f) Energy landscapes corresponding to the centroid of the parameter space. The relative *E_A_* values are shown. The ordinate axis is interrupted to indicate the unknown absolute height if the transition state.

By contrast, the interaction strength clearly increases for increasing *c*(MgCl_2_), green and blue in Fig. 3a. This observation is in agreement with the stiffening of vimentin networks in the presence of 5 mM MgCl_2_ at sufficiently long network formation times as shown in Fig. 1b. The interactions can reach several hundred pN, as shown in Fig. 2e, Fig. 3a and Fig. S18a, and are oftentimes even stronger than the optical trap. While interactions are stronger at low *c*(MgCl_2_) = 5mM than for the control condition, see Fig. 3a, the forceindependent binding rate *r_e,b_* is not increased, see Fig. 3b, indicating that interactions occur with the same probability. By contrast, at high *c*(MgCl_2_) interactions become more likely, see also Fig. S18a. The increase of *F_i_* for both *c*(MgCl_2_) measured here and the increase of *r_e,b_* solely at high *c*(MgCl_2_) explains why vimentin networks form “scaffold” like structures at *c*(MgCl_2_) < 10mM and collapse at high *c*(MgCl_2_) [16, 17]. We therefore hypothesize that it is a significant increase of the binding rate that causes the collapse of the networks and not the increased binding strength. This hypothesis is further supported, as we only observe zipping in the presence of 20 mM MgCl_2_, see Fig. S18d.

The strongest increase of binding rate, interaction strength and thereby also occurrence of zipping events is caused by the addition of high concentrations of MgCl_2_ and TX in combination (purple in Fig. 3a,b and Fig. S18). This “amplifying” effect of TX on MgCl_2_ is remarkable, as TX alone only weakly affects the interaction. It can possibly be explained by a swelling of the negatively charged and intrinsically disordered tail domains of the vimentin monomers, which decorate the filament surface, thus making them more accessible for interactions mediated by MgCl_2_. Because of this drastic change, we refrain from fitting the interactions in this condition with our simplified model.

#### A two-state model accurately describes network mechanics

From the interaction experiments, we extract the force-independent binding rate *r_e,b_*, Fig. 3b, and the distribution of interaction forces, Fig. 3a. For a quantitative comparison of the bonds formed in each condition, we model the interaction as a single bond. The filaments undergo force-dependent, stochastic transitions between the bound (*b*) and unbound (*u*) state that are described by Bell-Evans kinetics [29]. The binding (*r_b_*, see methods, Eq. (1)) and unbinding rates (*r_u_*, Eq. (2)) consist of a force-independent term (*r_e,b_* and *r_e,u_*, Eq. (3), Eq. (4)) that depends on the respective activation energies, *E_Ab_* or *E_Au_*, and the thermal energy *k_B_T*, and a force-dependent term that depends on *k_B_T*, the applied force, and the distance of the respective state to the transition state, *x_b_* or *x_u_*, see Fig. 3c [30]. The sum *x_c_* = *x_b_* + *x_u_* is constant due to detailed balance [31]. We calculate the distribution of *F_i_* for a constant *x_c_*, the *r_e,b_* derived from the experiment (Fig. 3b), and for pairs of x_u_ and *r_e,u_*, by simulating 1000 *F*(t) curves. Equivalent results are achieved by numerical solutions, as discussed in detail in Ref. [30]. Comparison of the experimental distribution and each simulated distribution based on the Kolmogorov-Smirnov test [32] with a 5% acceptance threshold, provides valid *x_u_* and *r_e,u_* pairs. The mean and standard deviations of the resulting distributions are shown in Fig. 3a. All parameter pairs for which the two-state model describes the experiment well, are plotted as pixels in Fig. 3d. The parameter spaces derived for the control and TX conditions overlap. The representative energy landscapes, calculated for the centroid of each respective parameter space, see circles in Fig. 3d (*r_e,u_*(control)=0.18 s^-1^, *r_e,u_*(0.16mM TX)=0.15s^-1^, *r_e,u_*(1.6mM TX)=0.20 s^-1^), shown in Fig. 3e, show no distinct differences. In the presence of MgCl_2_, *r_e,u_* decreases compared to the control (*r_e,u_*(5mM MgCl_2_)=0.08 s^-1^, *r_e,u_*(20mM MgCl_2_)=0.10s^-1^), which becomes apparent in a larger *E_A,u_* in Fig. 3f. These results are similar to values determined from rheometry on actin networks (0.43 ±0.06 s^-1^ [33]) but significantly larger than reported for vimentin networks (> 0.001s ^−1^ [8]) which is likely caused by surface effects in the macrorheology experiments.

The distance of the transition state to the bound state, *x_u_*, enters *r_u_* (Eq. (2)) in the factor that describes the force-dependent unbinding. Therefore, the decrease of both, *x_u_* and *r_e,u_* in the presence of MgCl_2_, leads to a mostly force-independent opening of the bond and thereby to high interaction forces. The presence of TX has the opposite effect, the parameter space reaches larger values of *x_u_*, indicating a more force sensitive bond between the filaments. The broader distribution of the parameters represented by the shaded areas in Fig. 3d, for the control condition and in the presence of TX are mostly due to the lower number of interactions.

These results support the assumptions we made during the analysis of the MPT experiments: As the filaments interact in all conditions, the resulting networks are not purely entangled but are connected by transient cross-links. However, in the control condition and in the presence of TX, the binding rate between filaments is low and interactions are weak, thus entanglements govern the network properties. In the presence of MgCl_2_, however, the unbinding rate is low, leading to stronger interactions, which in turn cause a more cross-linked network. As a low *c*(MgCl_2_) mostly affects the unbinding rate and not *x_u_*, *E_A,u_* can be increased without necessarily decreasing *E_A,b_*.

#### Conclusion

By combining experimental and modeling approaches, we are able to quantify three important contributions to biopolymer network properties and their temporal evolution, *i.e.* filament elongation kinetics, single filament mechanics and filament–filament interactions. We further determine which kinds of interactions, *i.e.* hydrophobic or electrostatic, are relevant for different processes involved in network formation and mechanics. We identify altered individual interactions between filaments as the cause of the network stiffening, where electrostatic effects play a key role. An increased interaction strength, caused by a less force-dependent unbinding, causes the stiffening of the networks that was previously reported in literature [6, 8–10], whereas hydrophobic interactions only play a minor role for filament interactions, but are, in the case of vimentin IFs, important during filament elongation. These results show that vimentin networks, even in standard buffer conditions without divalent ions, are not purely entangled but are connected by transient cross-links and the kinetics and strength of these transient cross-links strongly depend on divalent ions/electrostatic interactions [8, 10]. With further knowledge of the architecture of the network, in particular the number of potential interaction partners for each filament, the binding probability in the network may be determined in future studies. From our observations we conclude that cells need to explicitly suppress attractive interactions or even disassemble the filaments in a targeted manner, for example by phosphorylation [34], to re-organize the extremely stable cellular vimentin networks.

## 2 Methods

### Protein preparation

Human vimentin C328N with three additional amino acids, GGC, at the C-terminus is recombinantly expressed in *E.coli* and purified from inclusion bodies [21]. The protein is stored in denaturing conditions at −80 °C. The cysteines in vimentin dimers are labeled with Atto647N-maleimide (ATTO-Tec, Siegen, Germany) [35]. Before starting the assembly process, labeled and unlabeled protein is mixed to obtain a final labeling ratio of 4 %. The formation of tetramers is induced by reducing the urea concentration with 30 min dialysis steps at 6, 4, 2, 1, 0M urea in 2mM phosphate buffer at pH 7.5. After an additional over night dialysis step against 2mM phosphate buffer (PB) at pH 7.5 at 8°C, the protein concentration is determined by UV-Vis absorption spectroscopy. For single filament experiments, the protein is subsequently diluted to 0.2 g/L and assembled for further use: the assembly is triggered by dialysis against assembly buffer, 2 mM phosphate buffer containing 100mM KCl, pH 7.5, at 37°C over night. For rheology experiments, the protein is prepared in the same way with a labeling ratio of 0.9 % and is subsequently diluted to 1.2 g/L. Details of microrheology sample preparation are provided below.

### Stretching of single filaments

Filament stretching experiments are performed as described previously in Refs. [21, 24]. We note that for these experiments all filaments are assembled under control conditions and incubated in the respective measuring buffer (2 mM phosphate buffer containing 100 mM KCl, pH 7.5 with additional 0.16 mM TX or 5mM MgCl_2_) for 30 s prior to the stretching. Filaments directly assembled in the respective buffers are typically not long enough for these stretching experiments.

### Optical trap measurements of individual filament–filament interactions

Individual interaction measurements are performed with a high power optical trap (C-trap, Lumicks, Amsterdam, Netherlands) in quadruple trap (q-trap) mode. The C-trap set-up combines optical trapping with confocal microscopy with excitation lasers of wavelength 532 nm, and 638 nm and a pressure driven microfluidic flow cell. The four inlets of the microfluidic flow cell (I)-(IV), as shown in Fig. S1, are loaded with maleimide-functionalized polystyrene particles [21, 36] (4.5 *μ*m, Kisker Biotech, Steinfurt, Germany) in assembly buffer (I), pure assembly buffer (II), 1.3mg/L vimentin filaments in assembly buffer (III) and the respective measuring buffer (IV). The six different measuring buffers are: 2mM phosphate buffer with 100 mM KCl, pH 7.5 (assembly buffer) with (1) no addition, (2) 1.6mM (0.1%w/V) TritonX-100, (3) 0.16mM (0.01 %w/V) TritonX-100, (4) 5mM MgCl_2_, (5) 20mM MgCl_2_, or (6) 20mM MgCl_2_ and 1.6mM TritonX-100. In a microfluidic flow cell (see Fig. S16), single filaments are attached between two bead pairs and the binding is controlled by confocal microscopy. The filament between beads b3 and b4, filament f34, is rotated by 90° with respect to filament f12 between beads b1 and b2. The unstretched filaments are brought in contact in the measuring buffer. Filament f34 is moved across filament f12 in the same *x,y*-plane at 0.55 *μ*m/s and the force acting on b1 is recorded for further analysis. During the measurements, the positions of all particles are recorded by bright field imaging while the filaments are observed with confocal microscopy. If possible, the movement is stopped before bead b4 comes in contact with filament f12, the particles on f34 are lifted in *z*, and moved back to the starting position of the measurement. Contact between the filaments is established and another measurement is recorded. In the case of a very strong interaction, where filament f12 is not in the initial conformation when bead b4 reaches the y-position of f12, the movement is continued until the interaction ruptures or b4 comes into contact with f12. The category of “strong” interactions includes all interactions with *F_1y,max_* >100pN where the interaction is so strong that one of the beads is pulled out of the trap or one of the filaments ruptures. After failure of one of the filaments or a strong interaction, the particles are discarded.

### Analysis of the interaction data

The force acting on bead b1 in the direction of the movement of filament f34, *F*_1*y*,raw_, is plotted against the distance f34 has been moved, *d*, as shown in gray in Fig. S17a. As the particles are moved at a constant speed, the force-distance curves are equivalent to force-time curves. For each force-distance curve, regions without interaction are manually selected, as marked in blue in Fig. S17a. This is necessary as we observe a linear decrease of the force detected by trap1 over time, which can be attributed to the interaction of the trap potentials. The linear background in the regions of no interaction is subtracted from the raw data, black in Fig. S17a. At this stage, the force curves are classified into two groups: (i) all *F_1y_* (d) curves that do not have jumps exceeding 0.99 pN are considered ‘non-interactinG′, (ii) all other curves are sorted into the ‘interactinG′ class. For all curves in the ‘interactinG′ class, all data points exceeding the 2σ-limit of the non interaction region (blue) are marked as the interaction region (red). Single data points that exceed the 2σ-limit are removed manually. The duration that two filaments are interacting, as well as the total time of one experiment, are determined. There are three different types of interactions, as shown in Fig. 2c-e: i) ruptures, where *F_1y_* returns to 0pN, ii) *kinks,* i.e., ruptures where *F_1y_* does not decrease to 0pN and iii) strong interactions, i.e., cases where the experiment ends before a rupture is observed and *F_1y max_* >100 pN. Interactions where 0 < *F_1y_* < 100pN at the end of the experiment, i.e. without rupture, are excluded from further analysis as they cannot be sorted in any of the categories i)-iii). For all interactions, the force on the interaction site, *F*, is calculated with the geometry factor as described in the Supplementary Information and in Fig. S17b.

### Modeling single interactions

A detailed description of the model in provided in the Supplementary Information. In brief, the interactions between single vimentin filaments are modeled with a two-state model. The transitions between the bound (*b*) or an unbound (*u*) state are given by

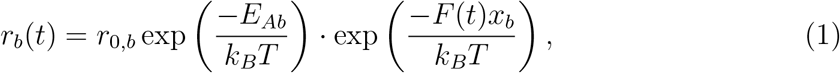

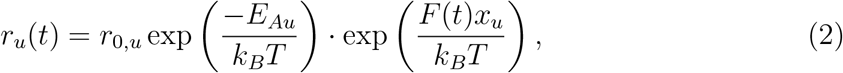

with the activation energy *E_Ab_* or *E_Au_*, the constant prefactor attempt rate *r_0,b_* or *r_0,u_*, the potential width *x_b_* or *x_u_*, and the thermal energy *k_B_T* at time *t*. The time dependent force *F(t)* includes the movement of the beads and the elastic and entropic stretching of the filaments, as sketched in Fig. S20. The transition rates have a force-independent contribution, *r_e,b_* or *r_e,u_*

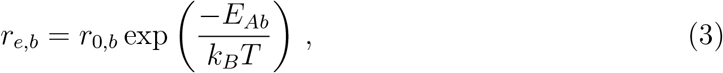

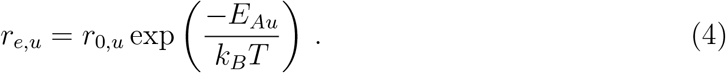

Force distributions, consistent with the experimental data, are generated by stochastic simulation and systematic variation of the unknown parameters.

### 2.1 Microrheology of vimentin networks

#### Microrheology of vimentin networks

##### Sample preparation

For assembly of vimentin networks, the final protein concentration is adjusted to 1 g/L. 6×buffers of 600mM KCl (assembly buffer), 600mM KCl with 30mM MgCl_22_, as well as 600 mM KCl with 0.96 mM TritonX-100, are prepared. Carboxylate-modified, green fluorescent microparticles of 2 *μ*m diameter (Molecular Probes, Eugene, Oregon, USA) are passivated by incubation with 0.1 g/L PLL(20)-g[3.5]-PEG(2 kDa) (SuSoS AG, Düben-dorf, Switzerland) at room temperature for 15 h. The microparticles are washed three times with PB before use [9]. Protein and buffer are mixed to to achieve the final protein concentration of 1 g/L protein in 1× buffer, and microparticles are added at less than 0.02% solids. Two no.1 cover slides (24 × 60 mm and 18×18mm, VWR, Radnor, Pennsylvania, USA) are cleaned by sonication in isopropanol and dried under a stream of N2_2_. For the sample chamber, a rectangular hole of 9×7mm is cut into 100-120 *μ*m thick doubled sided tape. The tape is placed on a clean 24× 60 mm cover slide, 6 μL of buffer with protein and microparticles are placed in the sample chamber and the device is sealed with a 18× 18 mm cover slide. The sample chambers are kept rotating at slow speed to avoid settling of the microparticles. After assembly in the dark at room temperature for the times given in Fig. 1, MPT measurements are performed.

##### Passive measurements

For passive microrheology measurements, we perform MPT using a commercial bright field microscope with a monochromatic LED with λ = 660 nm and a 60× objective (CFI Achromat FF, NA=0.80, WD=0.3mm, Nikon, Tokyo, Japan) with pre-implemented tracking software (Lumicks). We record a time series of the 2D microparticle position by tracking the diffraction ring patterns of microparticles with a quadrant interpolation algorithm. Traces are recorded for microparticles with a minimum distance of 30 *μ*m to all surfaces over the course of 6 min at a frame rate of 138 s^-1^.

##### Active measurements

We perform active microrheological experiments by oscillating the microparticle in the network with an optical trap (NanoTracker 2, JPK Instruments, Berlin, Germany) mounted on a Zeiss inverted optical microscope (AXIO Observer A1, Carl Zeiss, Jena, Germany). The setup is equipped with a water immersion objective (63×, C-Apochromat, NA = 1.2, Carl Zeiss) and an infrared laser (1024nm Nd:YAG, 5W). Trap stiffness and sensitivity are calibrated in assembly buffer from the power spectrum of the thermal fluctuations of several microparticles. We oscillate each microparticle with an optical trap at a laser power of 20 or 50 mW. The microparticle displacement and the phase shift of the oscillation of the particle with respect to the trap center are measured by back-focal-plane interferometry with a quadrant position detector.

##### Plateau modulus

To determine the frequency dependence of the storage modulus for each median G′ curve, we fit the curve in the frequency range where ω > 0.005s^-1^ and 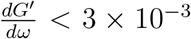 Pas by a power law *G′* = *C* exp(*p*_0_ω) using the least square method. The resulting fit curves and exponents *p*_0_ are presented in Fig. S12. We evaluate the storage modulus of the sample median curves at *ω* = 1 s^-1^ for the network initiation. We define the time point of the network initiation when the following requirements are first fulfilled: the filament length average *l_M_* stays constant, shown in Fig. 1c, and the MSD satisfies the condition max(MSD)< *a_particle_* = 2 *μ*m, as shown in Fig. 1b. From this time point on, the network mechanics vary because of network remodeling and not because of further entanglement due to filament elongation.

##### Relaxation time

The relaxation time *τ_e_ = 1/ω_e_*, where the mechanical response shifts from the deformation of the network *ω* < *ω_e_* to the deformation of the single filaments *ω* > *ω_e_* is extracted from the cross-over of the median curves of *G′* and *G″*. In a network, the bending of a single filament is constrained by other filaments. The relaxation time of the bending mode with the longest wavelength that is not constrained is *τ*e** [37].

##### Bundling parameter

At high frequencies, the shear modulus of networks of semiflexible filaments is given by [38]

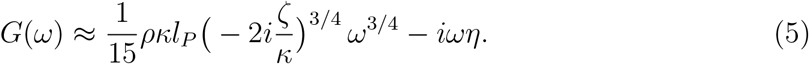

At these high frequencies, the response depends on the mass per length *ρ*, the persistence length *l_P_*, the lateral drag coefficient *ζ* and the bending stiffness *κ* = *k_B_Tl_P_*. We neglect the imaginary part [38] and fit our G″ curves at high frequencies, ω > 5s^−1^, by

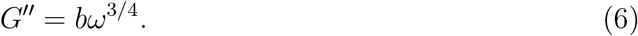

##### Imaging filament networks and single filaments

Confocal stacks of the samples used in microrheology experiments are recorded using an IX81 confocal microscope equipped with a 100× objective (Olympus, Hamburg, Germany). For each sample, the full height of the sample chamber is scanned with 5 *μ*m *z*-distance. Single filaments are imaged with wide-field fluorescence microscopy at 100 × magnification using an IX81 inverted microscope (Olympus). To follow the assembly speed of the filaments, vimentin protein and the respective assembly buffer are mixed at the same ratio as for microrheology experiments and diluted to 2 mg/L for imaging. The filaments are imaged on no.1 cover slides.

##### Analysis of filament lengths according to polymer step growth

We determine the lengths of the filaments in the fluorescence microscopy images, see Fig. S2, by manual tracing using ImageJ. We fit the resulting distribution with Flory’s theory of polymer step growth [11], using the ULF as the “monomer of the reaction” with a monomer length of *l_m_* = 43 nm [39].

## Acknowledgments

We thank J. Kraxner, and J.C. Thiele for helpful discussions, and S. Bauch for technical support.

This project was funded by the European Research Council (ERC) under the European Union’s Horizon 2020 research and innovation program (Consolidator Grant Agreement no. 724932). This research was conducted within the Max Planck School Matter to Life supported by the German Federal Ministry of Education and Research (BMBF) in collaboration with the Max Planck Society. The work further received financial support via an Excellence Fellowship of the International Max Planck Research School for Physics of Biological and Complex Systems (IMPRS PBCS) and the Studienstiftung des deutschen Volkes e.V.

## Author Contributions

S.Kö. conceived and supervised the project. A.V.S. performed and analyzed the single filament experiments. P.N. performed and analyzed active microrheology measurements. A.V.S. performed MPT measurements and P.N. and A.V.S. analyzed MPT experiments. A.V.S., P.N., S.Kö. and A.J. interpreted the microrheology results. C.L. and S.Kl. designed and performed numerical simulations. All authors contributed to writing the manuscript.

