## Supplementary information for "Multiscale mechanics and temporal evolution of vimentin intermediate filament networks"

[b] *Max Planck School “Matter to Life”*

[c] *Institute of Physical Chemistry, University of Göttingen, Tammannstr. 6, 37077 Göttingen, Germany*

[d] *Institute for the Dynamics of Complex Systems, University of Göttingen, Friedrich-Hund-Platz 1, 37077 Göttingen, Germany*

#### 1 Analysis of filament lengths according to polymer step growth

We determine the lengths of the filaments in the fluorescence microscopy images by manual tracing using ImageJ. The assembly of IFs occurs in two steps: combining lateral assembly to unit length filaments (ULFs) and longitudinal assembly by end-to-end annealing of the ULFs as shown in Fig. S1.

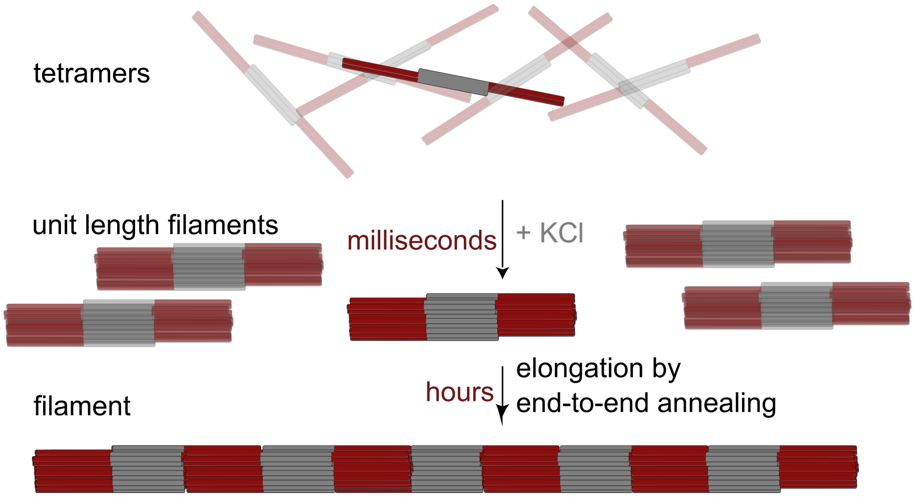


**Fig. S1 Filament assembly.** Sketch of the assembly process of vimentin IFs. ULFs form by lateral arrangement of eight tetramers. Filaments form by end-to-end annealing of the ULFs. Based on Ref. [1].

The longitudinal assembly is equivalent to a polymer step growth where the ULFs are the "monomers" of the polymerization. Following Flory's theory [2], the probability that a single reaction, the end-to-end annealing, has occurred is equal to $p$, the extent of the reaction. To build a polymer from $x$ monomers, $x-1$ reactions have to take place, which occur with the probability $p^{x-1}$. The probability to find an unreacted end group is $1-p$. The probability to find a polymer of length $x$ is therefore given by:

$$P_x = (1-p)p^{x-1} = \frac{N_x}{N}, \tag{S 1}$$

where $N$ is the total number of molecules of all sizes and $N_x$ is the number of molecules of length $x$. As we fit histograms of the length distributions, Fig. S3, with a certain bin width, we use the integrated form of equation (S 1) and optimize the fit for the full width of the bin by the least square method, the resulting $p$ values are shown in Fig. S4. To convert the lengths we measure here to the number of ULFs in one filaments, we use $l_m = 43$ nm [3], the repeat length of ULFs in IFs, as the "monomer size". We utilize the linear relation of weight and length, *i.e* a ULF has a constant weight and length, to calculate the length average $l_M$ from $p$, in analogy to the weight average:

$$l_M = \frac{l_m(1+p)}{1-p}. \tag{S 2}$$

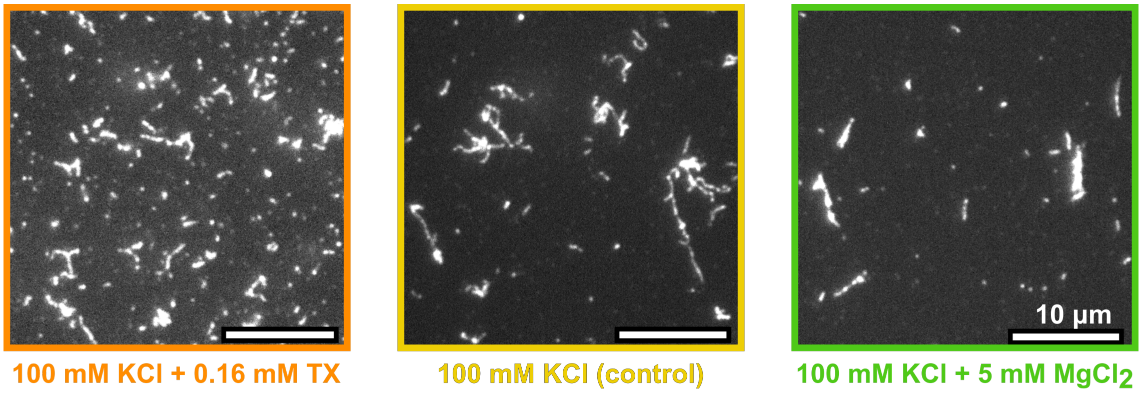


**Fig. S2 Example fluorescence microscopy images after 71 h assembly duration.** The filaments are assembled at 0.2 g/L in the respective buffer and diluted 100 fold for imaging. The corresponding length distributions of several of such images are shown in Fig. S3.

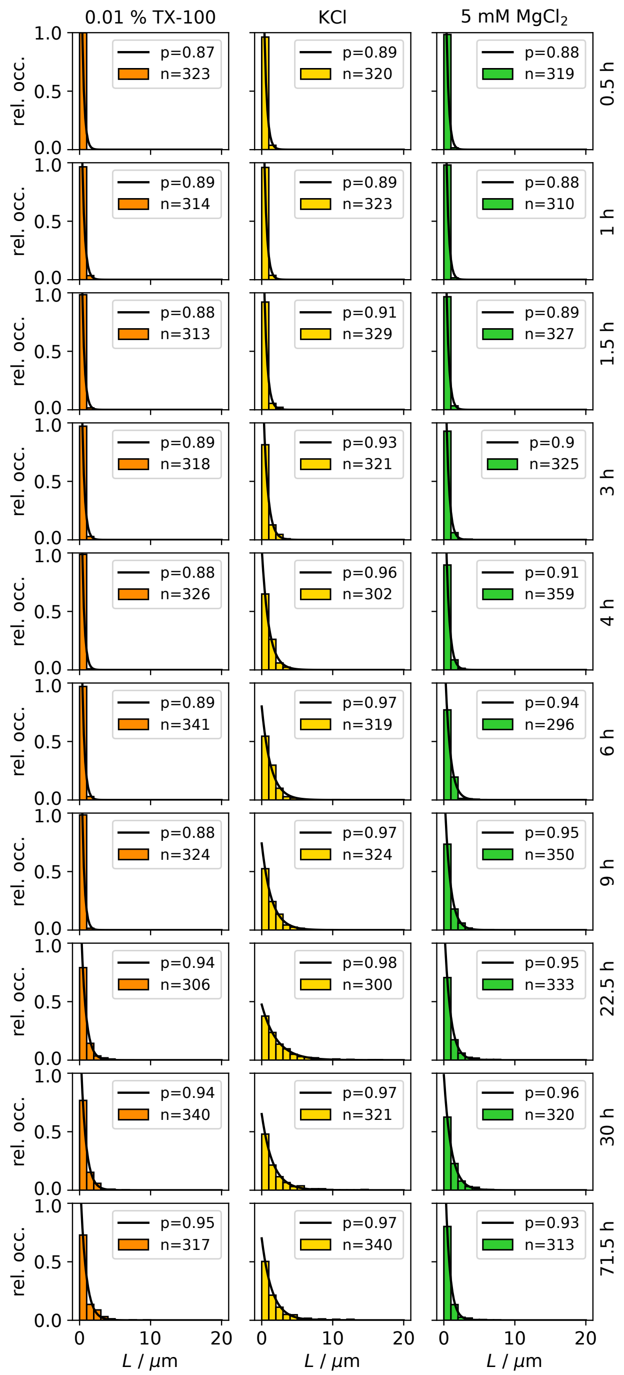


**Fig. S3 Histograms of filaments lengths measured by fluorescence microscopy images after the given assembly duration.** The filaments are assembled at 0.2 g/L and diluted 100 fold for imaging.

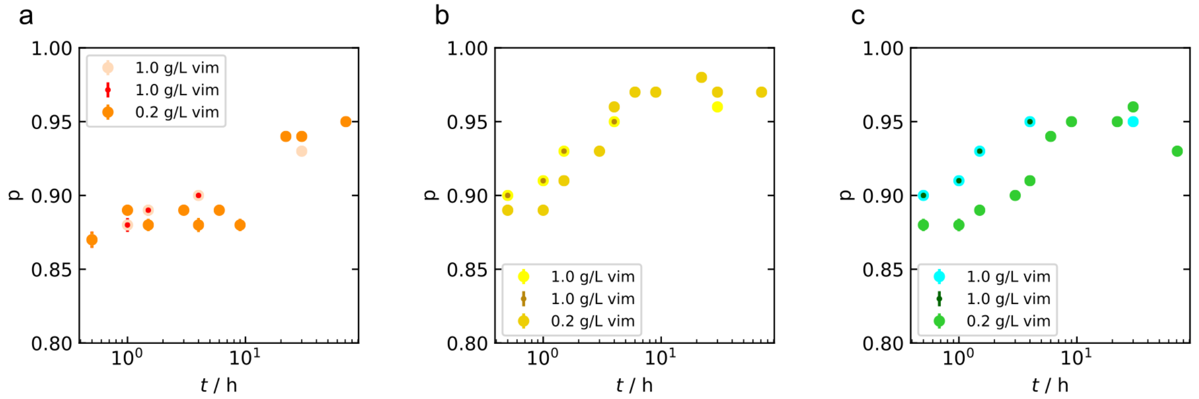


**Fig. S4 Extent of the filament elongation reaction.** The extent of the reaction, $p$, resulting from fits of the length histograms in Fig. S3, for protein concentrations of 0.2/L and 1.0 g/L. a) In assembly buffer + 0.16 mM TX, b) pure assembly buffer and c) in assembly buffer + 5 mM $MgCl_2$.

#### 2 Data analysis of passive microrheology measurements

The fluctuation-dissipation theorem can be applied to the recorded thermal motion of microparticles embedded in a viscoelastic medium that is in equilibrium on the time scale of the measurement. By utilizing the fluctuation-dissipation theorem, we imply that the medium, here the filament network, reacts to small, time-dependent external disturbances the same way it reacts to internal thermal fluctuations. The viscoelastic properties of a medium are characterized by the shear modulus $G^*(\omega)$, which is connected to the thermal fluctuations of the microparticles via a generalized Stokes-Einstein equation. A Laplace unilateral transformation – denoted by the tilde symbol – of the generalized Stokes-Einstein equation results in

$$\tilde{G}(s) = \frac{k_{\mathrm{B}}T}{\pi a s \left\langle \Delta\tilde{r}^2(s) \right\rangle} \tag{S 3}$$

with the particle diameter $a$ and the Laplace frequency $s$. The mean-squared displacement (MSD) $\left\langle \Delta r^2(\tau) \right\rangle$ is directly calculated from the thermal fluctuations of the microparticles for the lag time $\tau$. The difficulty lies in the Laplace transformation of the data over a limited frequency range, which can distort the result. To simplify the fitting process, a limited frequency range is described by a local power law and $G^*(\omega)$ determined analytically [4, 5]. The local power law is extracted from the logarithmic time derivative of the MSD.

$$\left|G^*\right| \approx \frac{k_{\mathrm{B}}T}{\pi a \left\langle \Delta r^2(1/\omega) \right\rangle \Gamma[1+\alpha(1/\omega)]} \tag{S 4}$$

$$\alpha(\tau) = \frac{d \ln \left\langle \Delta r^2(\tau) \right\rangle}{d \ln \tau} \tag{S 5}$$

Here, $\Gamma$ is the gamma function and $\alpha$ the local power-law exponent. Using the second order for this step improves performance of the algorithm [6]. We use a custom Matlab code based on this method to calculate $G^*$ [7]. Finally, the complex shear modulus $G^*$ is separated into the real part, the storage modulus $G'(\omega)$, and the imaginary part, the loss modulus $G''(\omega)$.

Interestingly, the heterogeneity observed in the confocal image of the network in Fig. 1a in the main text (bottom) is not obvious in the single MSD curves in Figs. S5-S7. We attribute this observation to the fact that only microparticles that are embedded in sufficiently dense network regions are accessible by our method and thus conclude that the regions where particles are embedded are homogeneous.

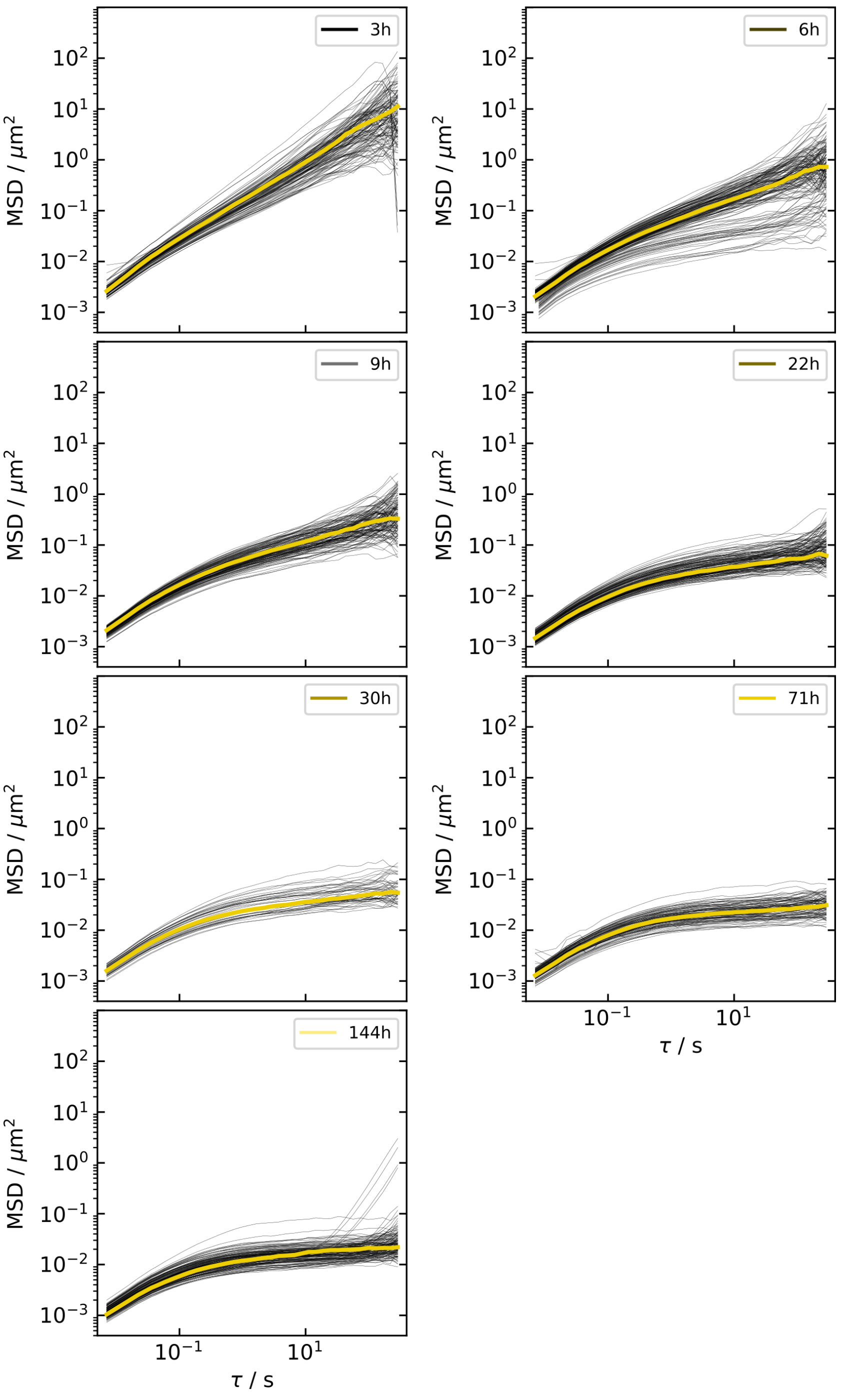


**Fig. S5 All single MSDs recorded for each assembly duration in KCl buffer.** The median curves are highlighted.

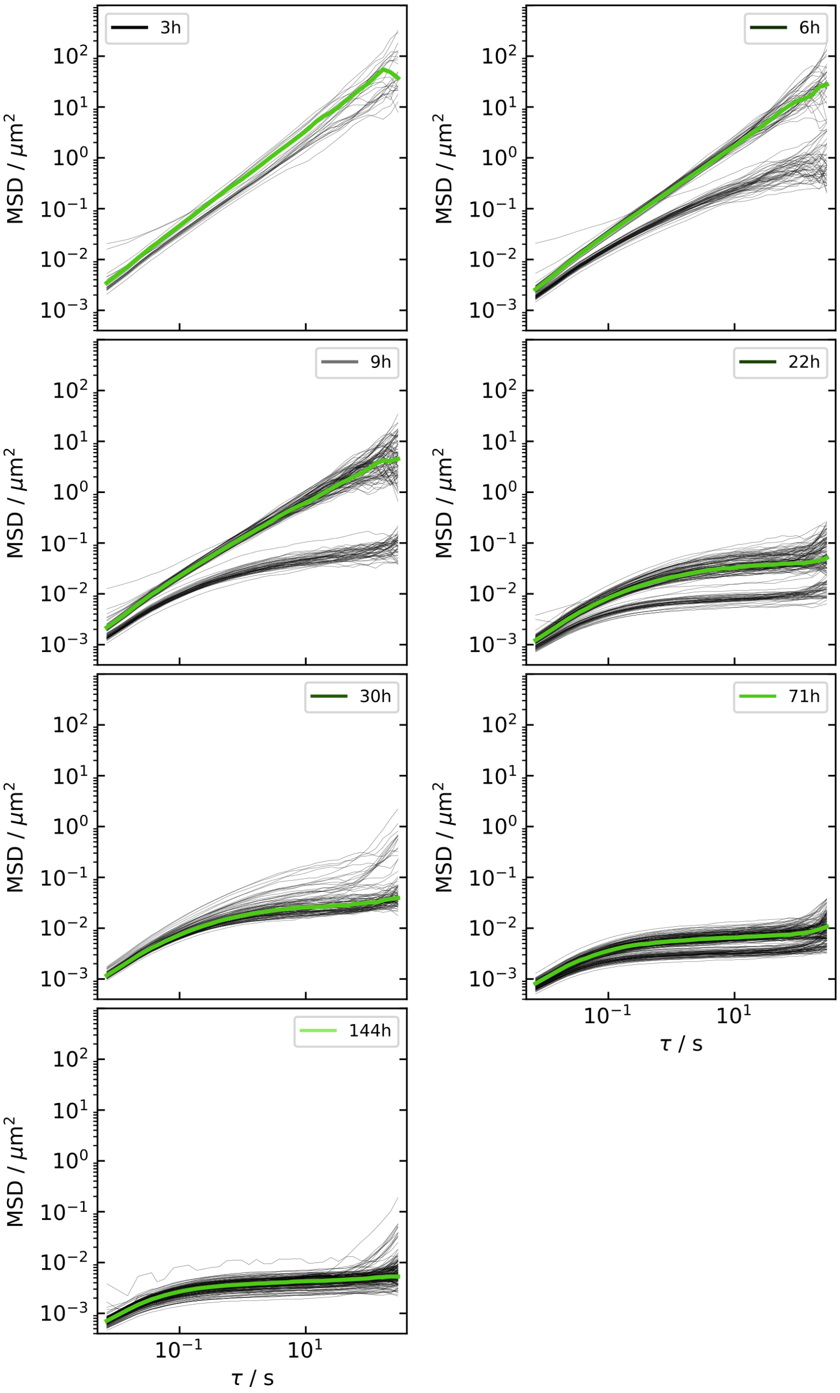


**Fig. S6 All single MSDs recorded for each assembly duration in 5 mM $MgCl_2$ buffer.** The median curves are highlighted. We observe variation between different samples prepared with the same concentration of $MgCl_2$, in particular at early time points. However, for each individual sample we obtain consistent MSD curves.

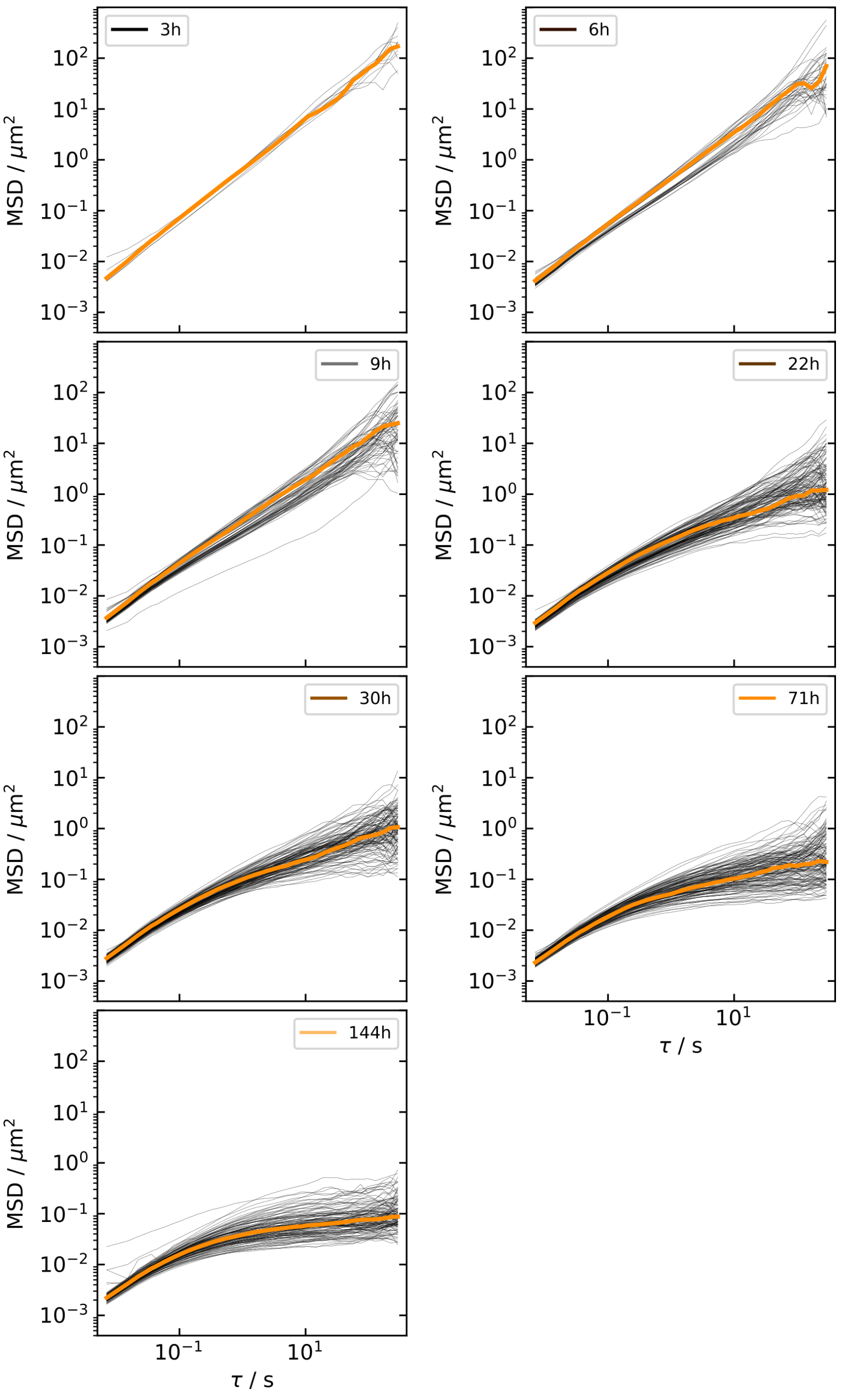


**Fig. S7 All single MSDs recorded for each assembly duration in 0.16 mM TX buffer.** The median curves are highlighted.

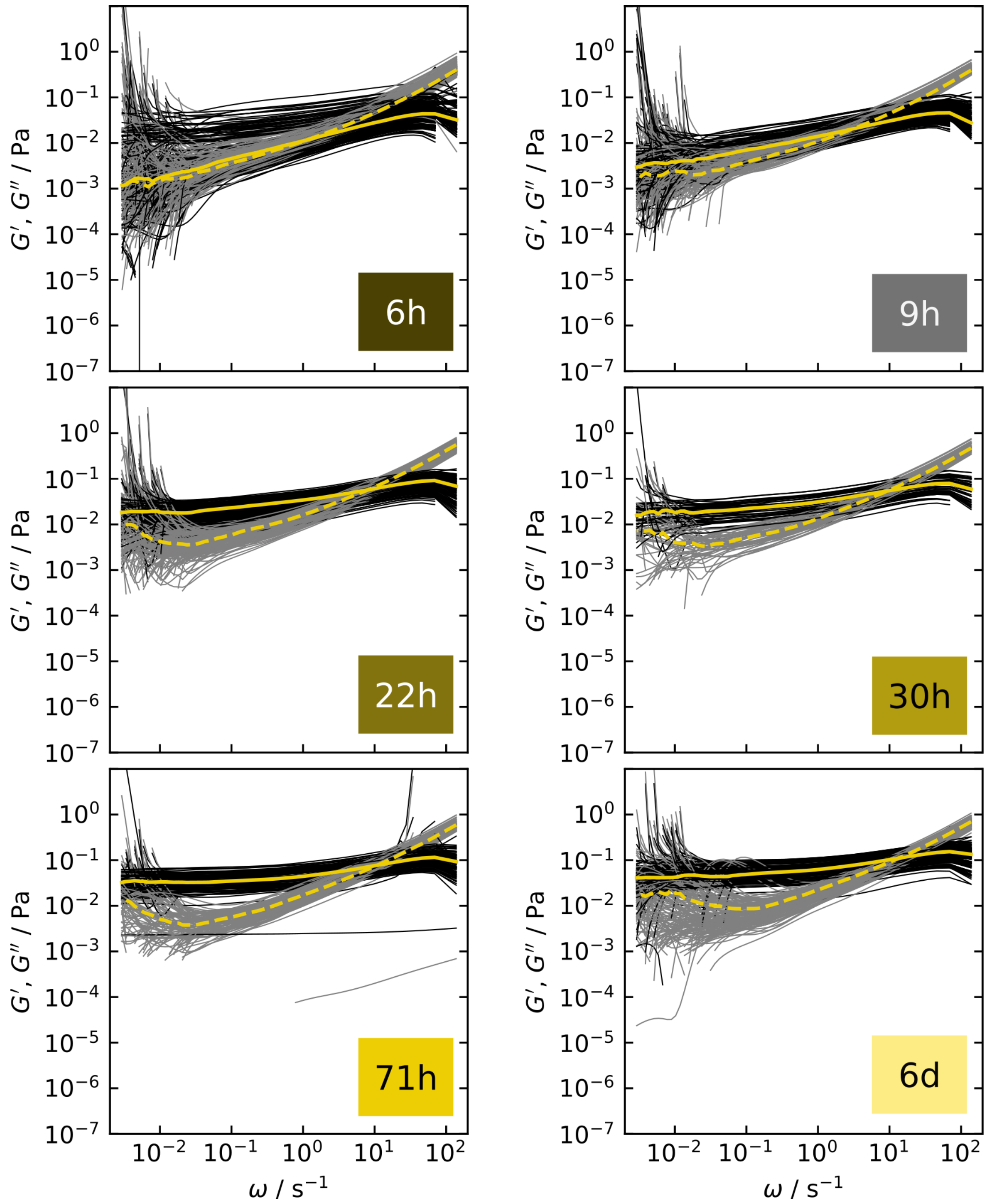


**Fig. S8 All single storage and loss moduli recorded for each assembly duration in KCl buffer.** The median curves are highlighted.

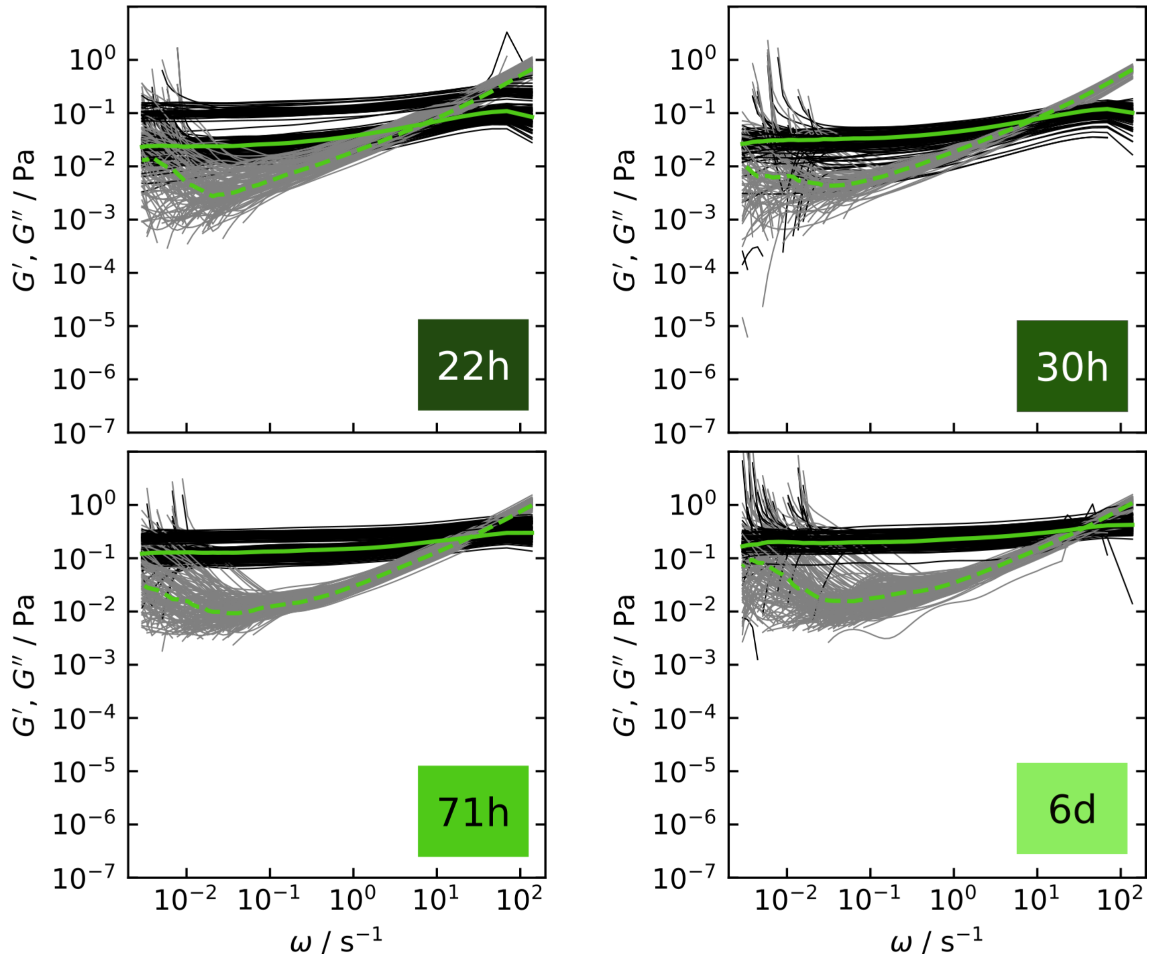


**Fig. S9 All single storage and loss moduli recorded for each assembly duration in 5 mM $MgCl_2$ buffer.** The median curves are highlighted.

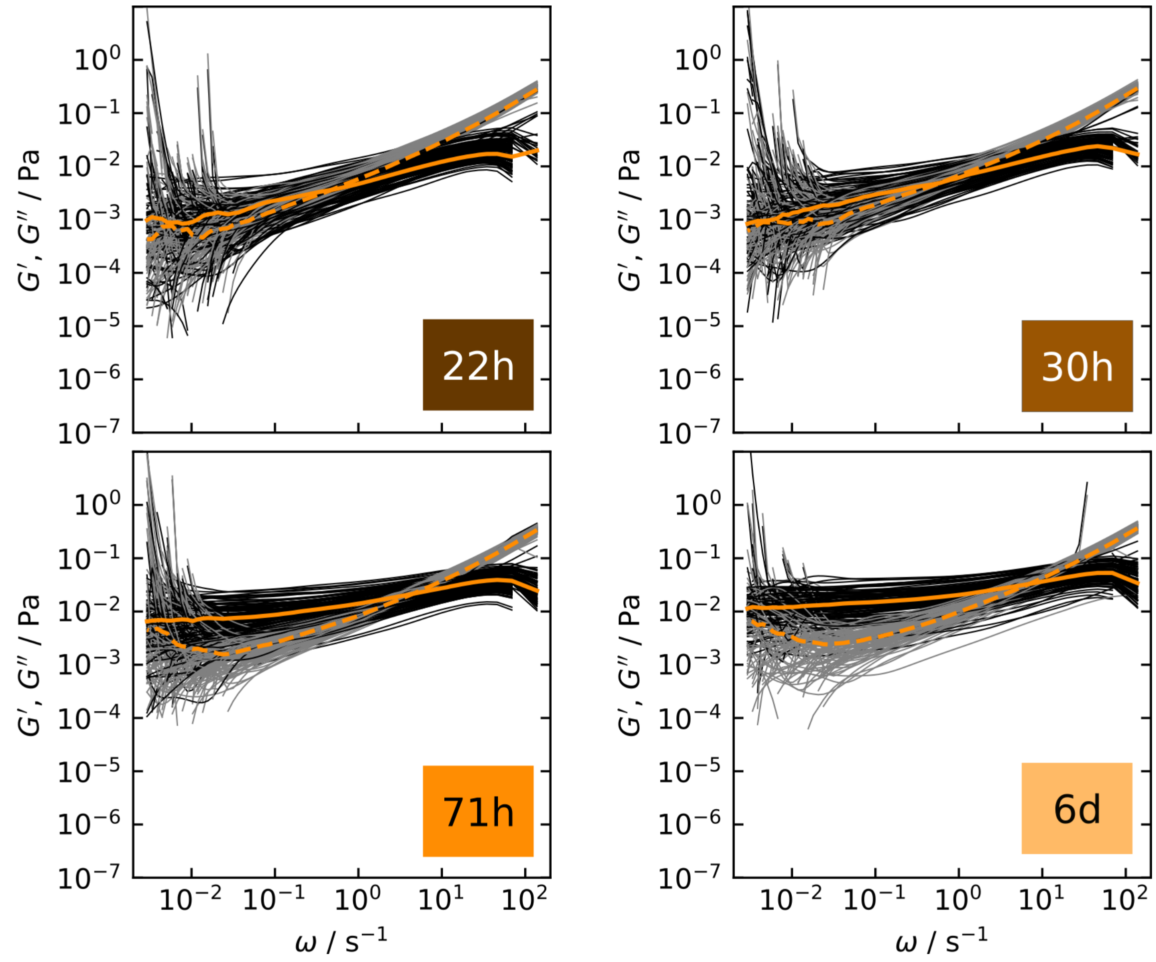


**Fig. S10 All single storage and loss moduli recorded for each assembly duration in 0.16 mM TX buffer.** The median curves are highlighted.

#### 3 Data analysis of active microrheology measurements

The amplitude-phase shift data are analyzed with a custom Matlab code. The particle is oscillated in the potential of an optical trap with the trap stiffness $\kappa$ and the amplitude $A$. The equation of motion for the particle is modeled with statistical random forces $f_R(t)$ acting on the particle and a drag force with the complex viscosity $\eta^*(\omega)$ as response of the surrounding medium:

$$m\ddot{x} = -6\pi a\eta^*(\omega)\dot{x} - f_R(t) + \kappa x + \kappa A e^{i\omega t}. \tag{S 6}$$

The equation is solved with a dampened and phase shifted sinusoidal motion:

$$x(t) = D(\omega)e^{i(\omega t - \delta(\omega))}, \tag{S 7}$$

where $D$ is the amplitude of the particle motion and $\delta$ its phase shift with regards to the driving force [8].

By neglecting the comparably small random forces and the inertial term for low oscillation frequencies, we can use a generalized Stokes-Einstein equation to connect the complex shear modulus $G^*(\omega)$ with $\eta^*(\omega)$

$$\eta^*(\omega) = \frac{G^*(\omega)}{\omega i}. \tag{S 8}$$

We arrive at an expression for $G^*$ with $D$ and $\delta$ as only remaining unknowns, which are determined by a sinusoidal fit to the trap motion as well as to the particle motion.

$$G^*(\omega) = \frac{\kappa}{6\pi a}\left(\frac{A}{D}\left(\cos(\delta(\omega) + i\sin(\delta(\omega)))\right) - 1\right) \tag{S 9}$$

This complex term is separated into the real part, the storage modulus $G'(\omega)$, and the imaginary part, the loss modulus $G''(\omega)$ [9].

$$G'(\omega) = \frac{\kappa}{6\pi a}\left(\frac{A}{D}(\cos(\delta(\omega))) - 1\right) \tag{S 10}$$

$$G''(\omega) = \frac{\kappa}{6\pi a}\left(\frac{A}{D}(\sin(\delta(\omega)))\right). \tag{S 11}$$

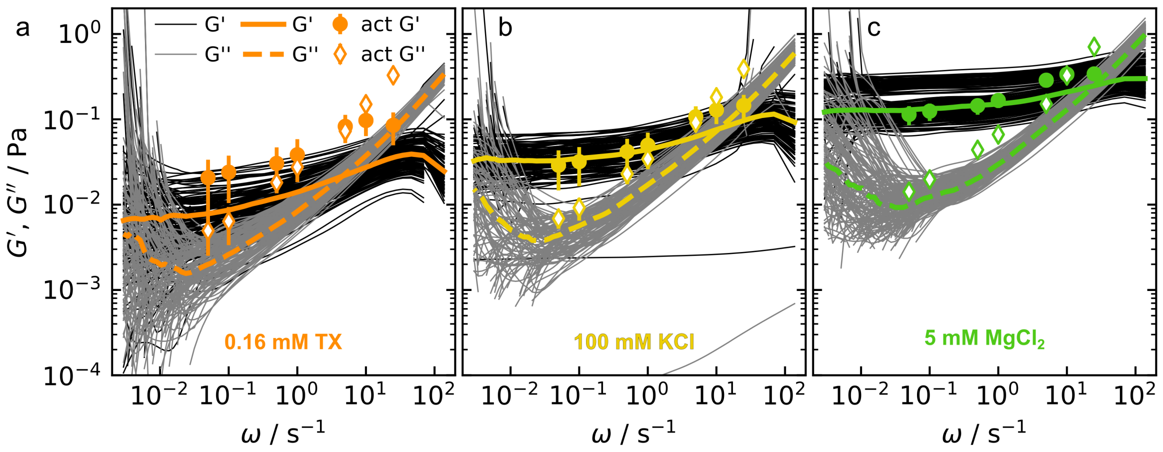


**Fig. S11 Comparison of active and passive rheology.** The single $G'$ and $G''$ curves from passive measurements after 71 h assembly time are shown by thin black and gray lines, respectively. The median of the samples is plotted with a bold line ($G'$) and bold, dashed line ($G''$). The active measurements are recorded on the same day. Rheological properties for measurements in 100 mM KCl plus a) 0.16 mM TritonX-100, b) nothing or c) 5 mM $MgCl_2$.

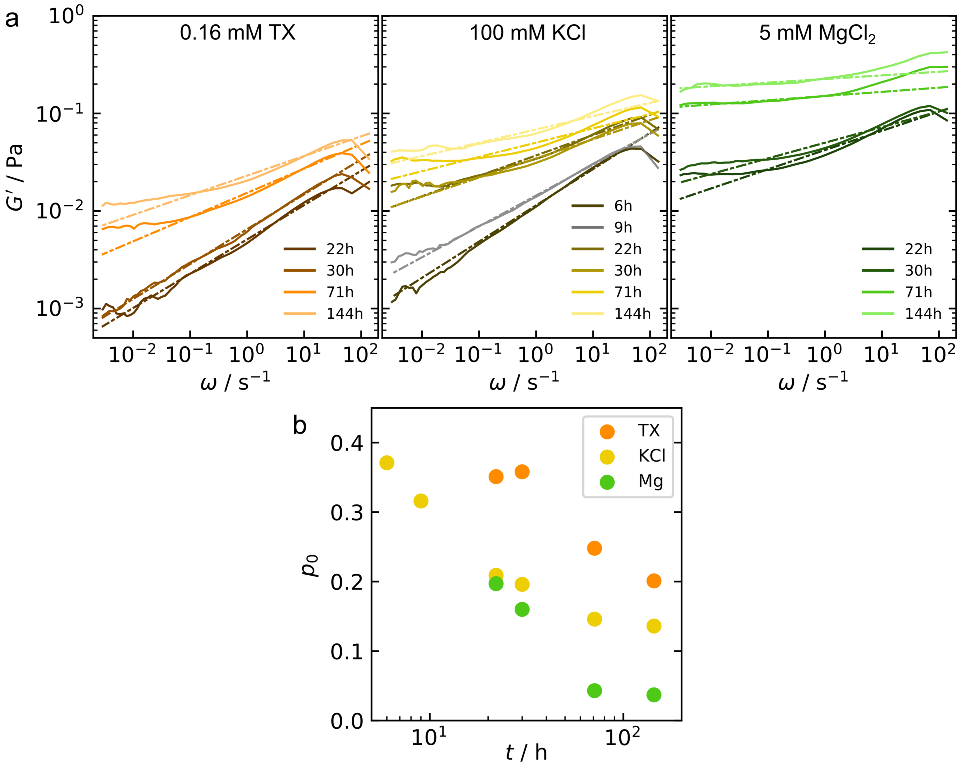


**Fig. S12 Frequency dependence of the storage modulus.** a) The exponent $p_0$ of a power law fit to the storage modulus $G'$ describes the frequency dependence of the storage modulus. The median $G'$ curves of the samples (solid line) and the resulting fit (dashed line) are shown. b) Comparison of the exponents of the power law fits over the network assembly duration.

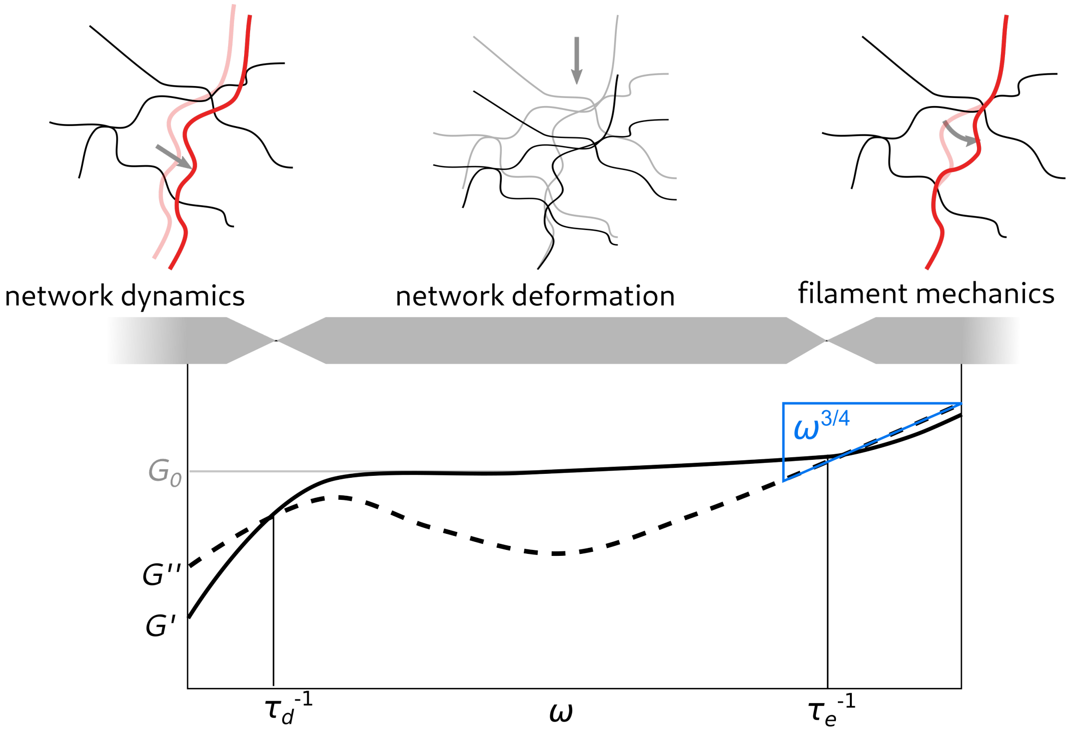


**Fig. S13 Three regimes in the viscoelastic properties of semiflexible polymer networks.** At low frequencies $\omega < \tau_d^{-1}$ the response is due to flow of the network by self-diffusion of the filaments. At intermediate frequencies, between the disentanglement time and the relaxation time, $\tau_d^{-1} > \omega > \tau_d^{-1}$, the network deformation is probed. The slope of the plateau depends on the entanglement or cross-linking of the network. At high frequencies $\omega > \tau_e^{-1}$, the mechanics, *i.e.* bending, of single filaments or bundles dominate the response.

#### 4 Relative bundling parameter

For comparison of the bundling parameter $b$ (Fig. 1g in the main text) between the samples at the different time points, we define $b(9h, \mathrm{KCl})$ as the reference value for a network without bundles, $b_f$.

$$G(\omega) \approx \frac{1}{15}\rho\kappa l_P\big(-2i\frac{\zeta}{\kappa}\big)^{3/4}\,\omega^{3/4} - i\omega\eta. \tag{S 12}$$

At high frequencies, the response depends on the mass per length $\rho$, the persistence length $l_P$, the lateral drag coefficient $\zeta$ and the bending stiffness $\kappa$ .

We fit the $G''$ curves at high frequencies, $\omega > 5\ \mathrm{s}^{-1}$, by

$$G'' = b\omega^{3/4}. \tag{S 13}$$

From equation (S 12) and $\kappa = k_B T l_P$ we obtain

$$b = \frac{1}{15}\rho k_B T l_P^2\big(-2i\frac{\zeta}{l_P k_B T}\big)^{3/4}. \tag{S 14}$$

In equation (S 14), $\rho$, $l_P$ and $\zeta$ depend on the number of filaments per bundle, $N$, as described in the following. The mass per length is additive, $\rho_b = N\rho_f$. The lateral drag coefficient is defined as

$$\zeta = \frac{4\pi\eta}{\ln(0.6\lambda/d)} \tag{S 15}$$

with the $\eta$ solvent viscosity, the filament or bundle diameter $d$ and the characteristic length scale $\lambda$, which can be the mesh size $\xi$ [10]. The dependence of the persistence length on $N$ in worm-like bundles depends on the coupling of the filaments within the bundle. In the case that the filaments are uncoupled, the persistence length of the bundle $l_{Pb}$ depends linearly on the persistence length of the single filaments $l_{Pf}$ [11]:

$$l_{Pb} = N l_{Pf}. \tag{S 16}$$

Using this assumption, we obtain

$$b_{rel} = \frac{b}{b_f} = \left(\frac{\zeta_b}{\zeta_f}\right)^{3/4} N^{9/4}. \tag{S 17}$$

For fully coupled bundles, we obtain [11]:

$$l_{Pb} = N^2 l_{Pf}, \tag{S 18}$$

and therefore

$$b_{rel} = \frac{b}{b_f} = \left(\frac{\zeta_b}{\zeta_f}\right)^{3/4} N^{17/4}. \tag{S 19}$$

The lateral drag coefficients $\zeta_f$ for filaments and $\zeta_b$ for bundles have a negligible effect in this relation. In Fig. S14f, the relative bundling parameter is plotted against $N$. These calculations allow for a rough estimate of the average number of filaments per bundle.

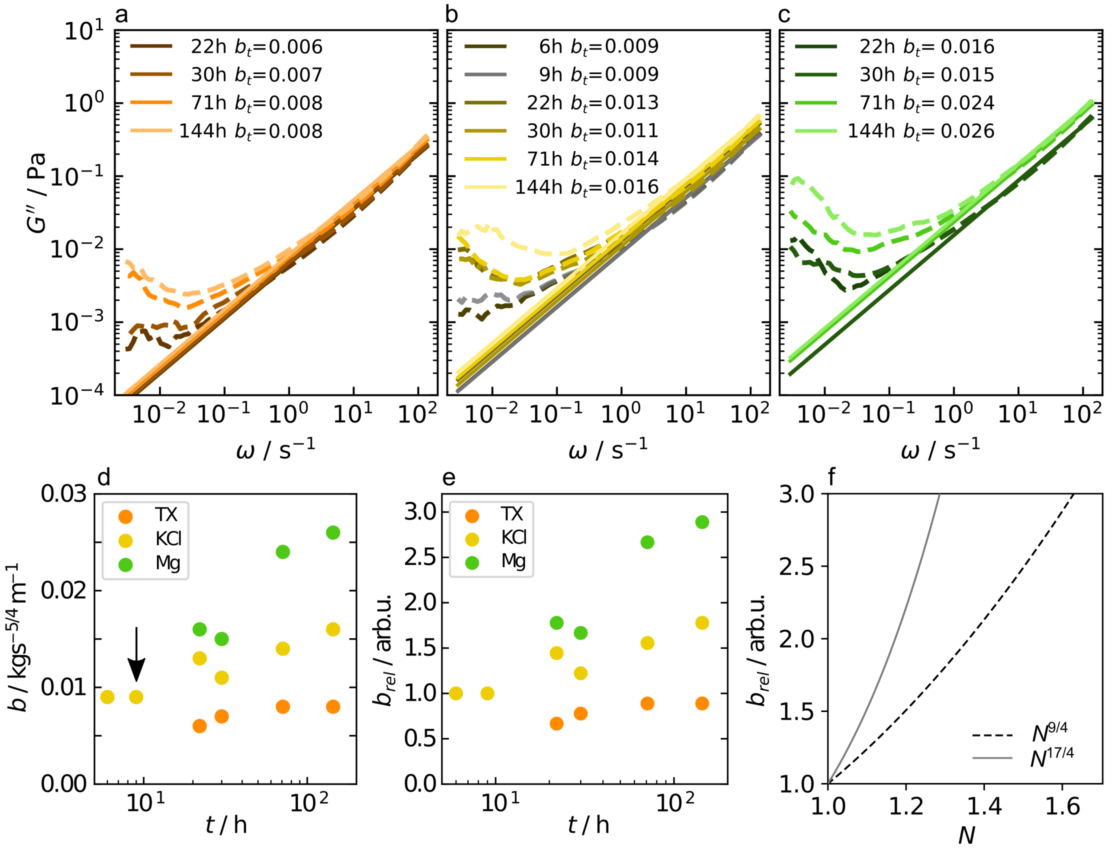


**Fig. S14 Relative bundling parameter.** a) Fit of $G'' \propto b\omega^{3/4}$ to the data involving TX, b) only KCl, and c) $MgCl_2$. d) Resulting fit parameter $b$ plotted against the assembly time of the network. e) The bundling parameter $b_{rel}$, relative to the value obtained for initial networks state of the the KCl sample after 9 h, marked with the arrow in d). f) Dependence of $b_{rel}$ on the average number of filaments per bundle. For fully coupled bundles $b_{rel} \propto (\frac{\zeta_b}{\zeta_f})^{3/4} N^{17/4}$, solid line, and for uncoupled bundles $b_{rel} \propto (\frac{\zeta_b}{\zeta_f})^{3/4} N^{9/4}$, dashed line. The ratio of the lateral drag coefficients $(\frac{\zeta_b}{\zeta_f})^{3/4}$ only weakly influences $b_{rel}$ and is neglected here.

### 5 Single filament stretching

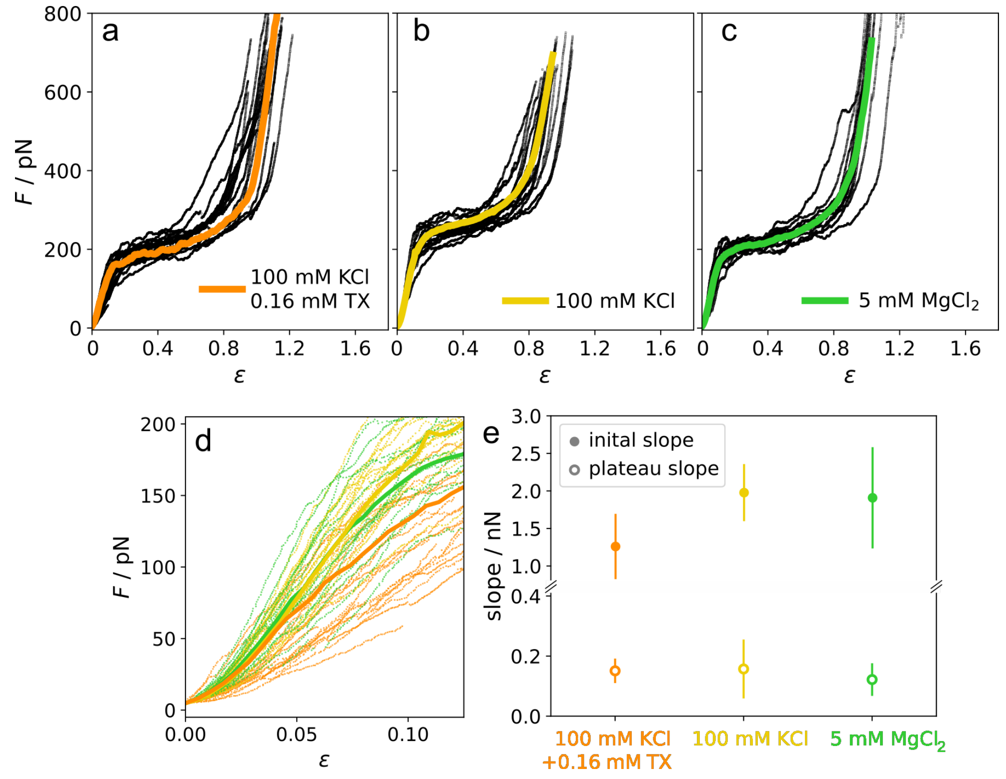


**Fig. S15 Stretching single filaments.** a) Force strain curves recorded for single vimentin filament stretching in the optical trap. The filaments are assembled in assembly buffer and measured after 30 s incubation in assembly buffer + 0.16 mM TritonX-100 (TX), b) assembly buffer, c) 2 mM phosphate buffer + 5 mM $MgCl_2$ at pH 7.5. The data shown in b) and c) are published in Ref. [12]. The single curves are shown in black and the average curve in the respective color. d) Comparison of the initial, linear increase of the force-strain curves for all three conditions, showing the slight softening of the filaments in TX. e) Comparison of the slope of a linear regression for the initial regime and the plateau regime. We show the mean of all single curves and the standard deviation (error bars). The slope of the plateau is constant for all conditions whereas the initial slope is slightly lower in TX than in assembly buffer and Mg-buffer.

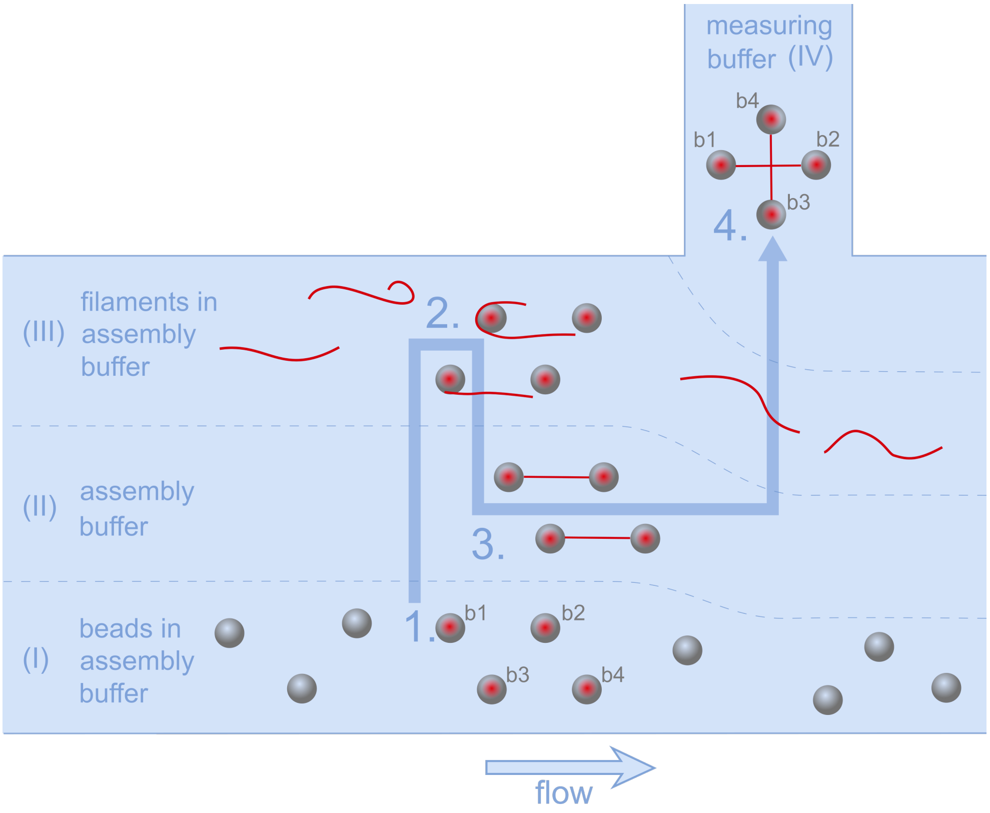


**Fig. S16 Schematic of the experimental protocol for single interaction measurements in the optical trap.** Laminar flow enables separation of substances in channels (I)-(IV). Four beads (b1-b4) are captured in the optical trap at position 1. in channel (I). The beads are calibrated in channel (II) and subsequently moved to position 2. in channel (III) to capture vimentin filaments. The beads are moved to position 3. in the buffer channel (II) to ensure a single filament is connected to the beads pair at both ends. After rotating one bead pair at position 3., the filaments are moved to position 4. in the measuring buffer in channel (IV). The filaments are brought into contact at position 4., where measurements are performed.

The force acting on the interaction site is

$$F = F_{1y} + F_{2y}. \tag{S 20}$$

However, in our setup, we can only directly measure forces acting on bead b1 and not the forces acting on bead b2. We therefore calculate $F$ from the position of filament f34 on filament f12. The forces in $x$-direction have to be balanced, so $F_{1x} = F_{2x}$. With the angle $\alpha_i$, $(i = [1, 2])$ between the deflected and non-deflected filament section,

$$\tan\alpha_1 = \frac{F_{1y}}{F_{1x}} = \frac{\Delta y_i}{l_1 + a/2} \qquad \text{and} \qquad \tan\alpha_2 = \frac{F_{2y}}{F_{2x}} = \frac{\Delta y_i}{l_2 + a/2}, \tag{S 21}$$

$F_{2y}$ can be expressed in terms of $F_{1y}$

$$F_{2y} = F_{1y}\frac{l_1 + a/2}{l_2 + a/2}. \tag{S 22}$$

By using $l_1 + l_2 = l_{12}$ and equation (S 20), the interaction force $F$ is

$$F = F_{1y}\left(\frac{l_{12} + a}{l_2 + a/2}\right). \tag{S 23}$$

All measurements that show interactions are corrected by their respective geometry factor to obtain the actual interaction force $F$, shown in Figs. 2 and 3 in the main text, from the measured force $F_{1y}$. The maximum force $F_i$, reached before a rupture or kink, is plotted in Fig. 3a in the main text.

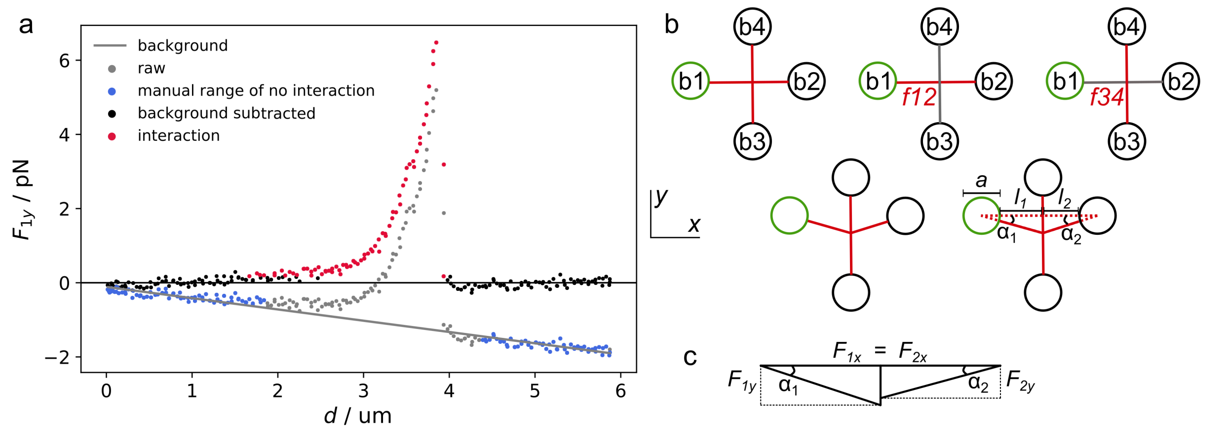


**Fig. S17 Analysis of single interaction experiments.** a) The force $F_{1y}$ is plotted against the relative position change $d$ of bead pair f34. The raw data , $F_{1y,\text{raw}}$, are plotted in gray. Two regions without interaction are chosen manually (blue) and a linear regression is calculated for those data points (gray line). This line is subtracted from the raw data to obtain the corrected force $F_{1y}$. The standard deviation of the regions without interaction (black) is calculated and all data points exceeding the $2\sigma$-limit of this region are classified as interactions, marked in red. b) To obtain the interaction force $F$, the background corrected forces are further multiplied by a geometry correction factor to account for filament f34 being off-center on filament f12 as described in equation (4) in the main text. c) Force diagram for the geometry shown in b).

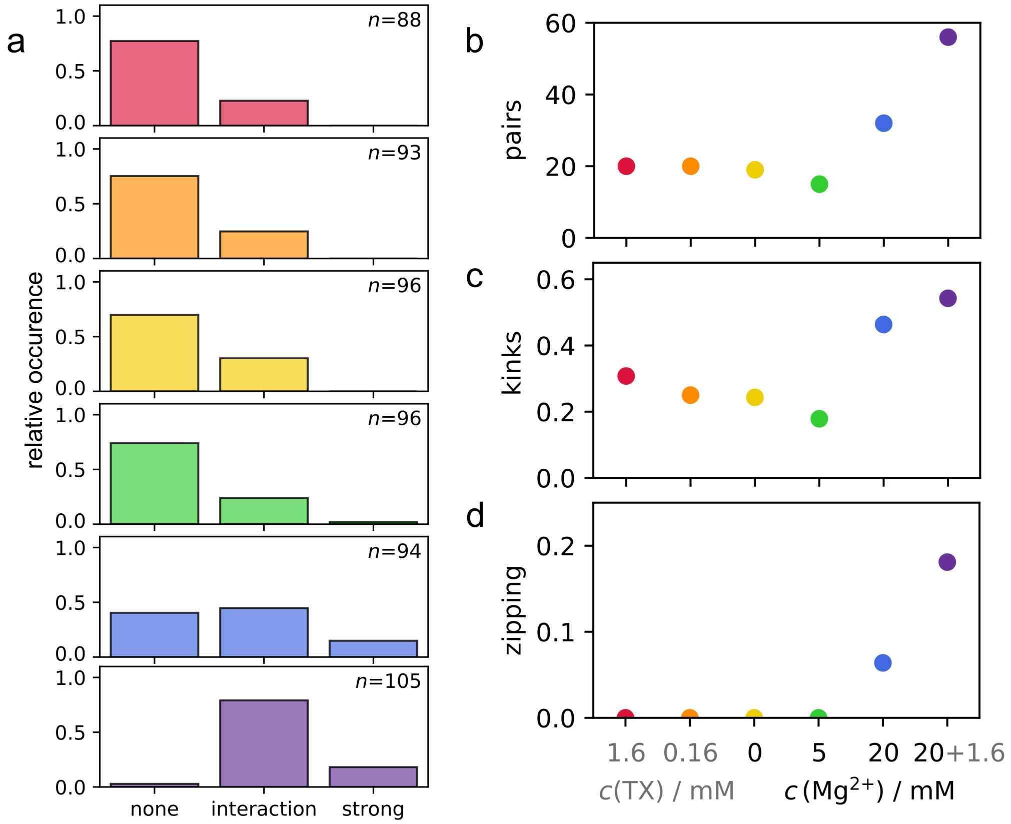


**Fig. S18 Interactions types.** a) The relative occurrence of measurements without any interaction, measurements with only interactions that show the characteristic force jump of a breaking interaction, and strong interactions, where the measurement ends before a force jump is observed. The strongest interaction of each measurement defines the category. $n$ denotes the number of measurements. b) The number of filament pairs measured in each condition. Pairs, where filaments are stable and interaction strengths do not exceed the maximal trap forces can be measured multiple times. c) The relative occurrence of kinks in all, *i.e.* normal and strong, interactions. d) Fraction of zipping events in all measurements. Color code as in main text.

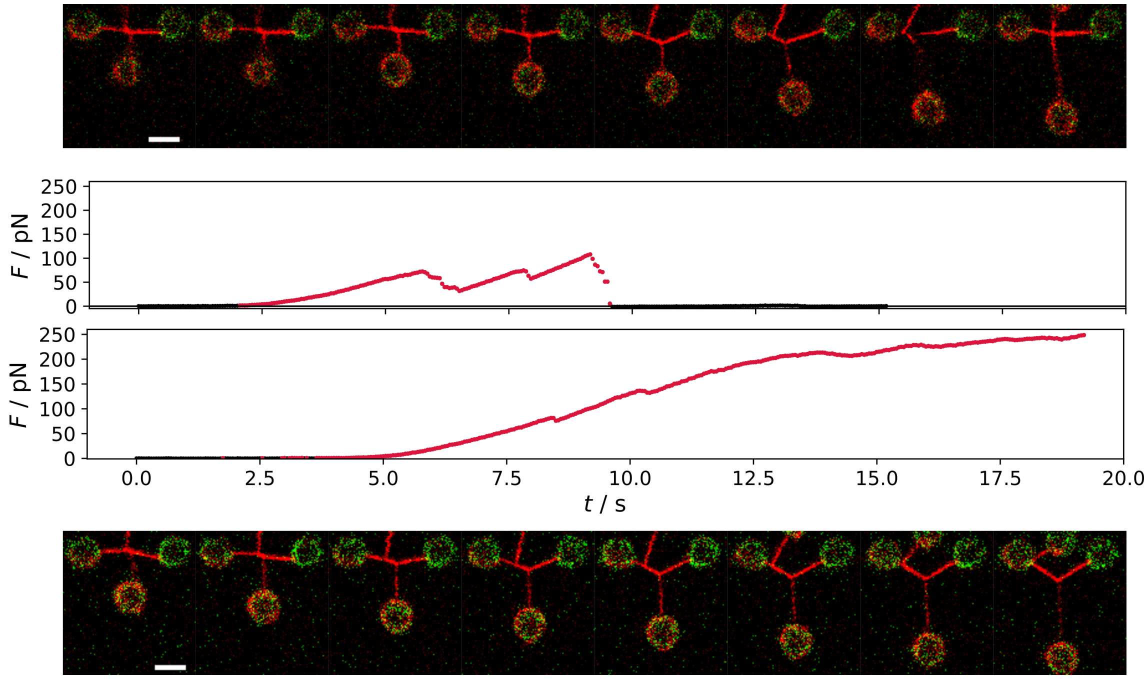


**Fig. S19 Zipping during interaction experiments.** Confocal scans and respective force curves for unzipping (top) and zipping (bottom). The confocal videos are available as Supplementary movies S1 and S2. The scale bar is 5 $\mu$m.

#### 7 Modeling single interactions

##### 7.1 Reaction rates

We model single interactions between two vimentin intermediate filaments with a two-state model since the filaments can be either in a bound ($b$) or an unbound ($u$) state. We describe the transition between the two states, *i.e.* the binding and unbinding rate, $r_b$ and $r_u$, respectively, with Bell-Evans kinetics [13]. The reaction rates, with the force $F(t)$, the activation energy $E_{Ab}$ or $E_{Au}$, the constant prefactor attempt rate $r_{0,b}$ or $r_{0,u}$, the potential width $x_b$ or $x_u$ and the thermal energy $k_BT$ at time $t$ are

$$r_b(t) = r_{0,b} \exp\left(\frac{-E_{Ab}}{k_BT}\right) \cdot \exp\left(\frac{-F(t)x_b}{k_BT}\right), \quad \text{(S 24)}$$

$$r_u(t) = r_{0,u} \exp\left(\frac{-E_{Au}}{k_BT}\right) \cdot \exp\left(\frac{F(t)x_u}{k_BT}\right). \quad \text{(S 25)}$$

The transition rates have a force-independent contribution, $r_{e,b}$ or $r_{e,u}$

$$r_{e,b} = r_{0,b} \exp\left(\frac{-E_{Ab}}{k_BT}\right), \quad \text{(S 26)}$$

$$r_{e,u} = r_{0,u} \exp\left(\frac{-E_{Au}}{k_BT}\right), \quad \text{(S 27)}$$

and a force-dependent term, scaling with the distance to transition state $x_b$ or $x_u$. The sum of $x_b$ and $x_u$ has a constant value $x_c$ to ensure detailed balance [14].

The rate equations may be either solved by simulating many single interaction events or numerically, as described in detail in Ref. [15].

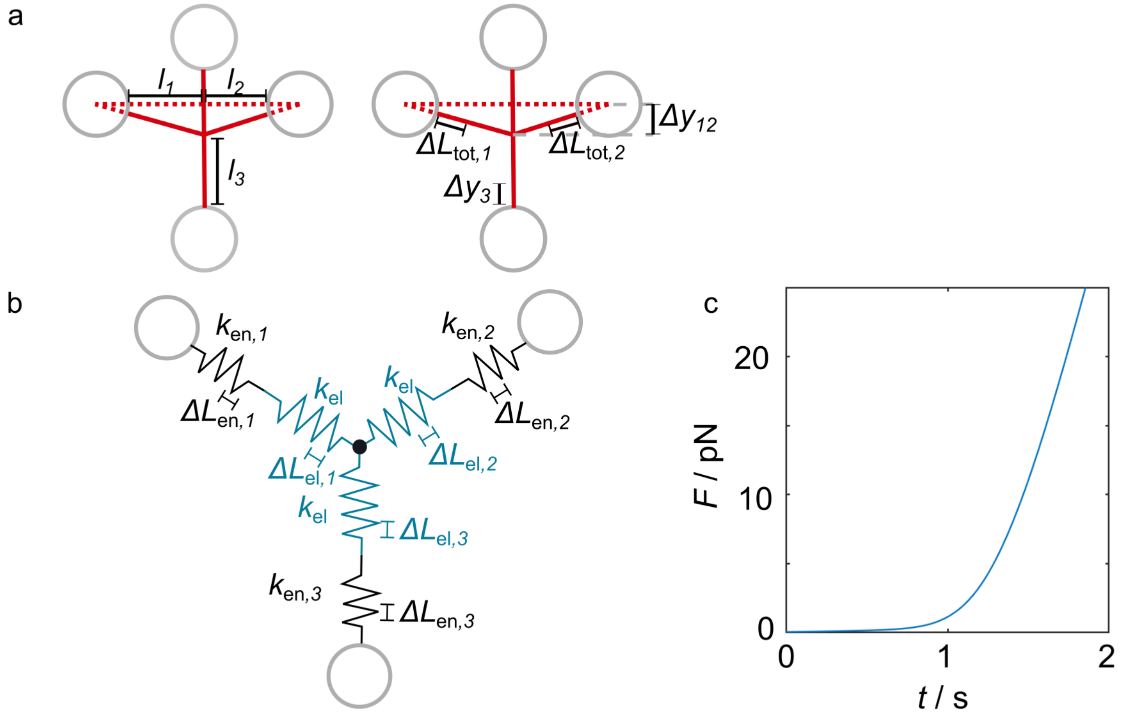


**Fig. S20** Parameters required for the microscopic model qualitatively describing the force increase in the quadruple optical tweezers experiment. a) Lengths and extensions of the filaments. b) Microscopic model of two interacting filaments with the spring constants required to calculate the force increase on the interaction. c) Theoretically expected force increase resulting from the microscopic model.

#### 7.2 Microscopic model of the filament interaction

To solve equations (S 24) and (S 25), we derive the functional shape of the force increase $F(t)$ with a microscopic modeling approach. To this end, we describe the filaments as a combination of springs. Two factors contribute to the force increase at the interaction site: 1. The elastic stretching and the entropic stiffening of the vimentin filaments, and 2. the variation of the deflection angle between the filament sections of filament f12 and the direction of motion of beads b3 and b4.

##### 1. Elastic and entropic filament stretching

We model vimentin filaments with a spring constant $k_{f,i}$ as consisting of an elastic

($k_{\mathrm{el}}$), and an extension-dependent, entropic ($k_{\mathrm{en},i}$) spring that are connected in series, as sketched in Fig. S20b, where $i$ is the index for the $i^{\mathrm{th}}$ filament section as marked in Fig. S20a:

$$k_{f,i} = \left(\frac{1}{k_{\mathrm{el}}} + \frac{1}{k_{\mathrm{en},i}}\right)^{-1} . \tag{S 28}$$

The elastic spring constant of the of vimentin filaments is set to $k_{\mathrm{el}} = (0.22 \pm 0.06)$ pN/nm [16]. To calculate the entropic spring constant, we assume a worm-like chain model to describe the end-to-end extension $l_{ee,i}$ of the $i^{\mathrm{th}}$ filament section, with the entropic force $F_e$, the persistence length $l_P$, $k_BT$ and the contour length of the $i^{\mathrm{th}}$ filament section $l_{c,i}$ [17]:

$$\frac{F_e l_P}{k_BT} = \frac{l_{ee,i}}{l_{c,i}} + \frac{1}{4\left(1 - \frac{l_{ee,i}}{l_{c,i}}\right)} - \frac{1}{4} . \tag{S 29}$$

We set the initial stretch $c_s$ of the filament to $c_s = l_{ees,i}/l_{c,i}$, where $l_{ees,i}$ is the initial end-to-end extension. With the extension of the entropic spring $L_{\mathrm{en},i}$, the end-to-end distance is $l_{ee,i} = l_{ees,i} + L_{\mathrm{en},i}$. We approximate the entropic spring constant of a filament section by the derivative of the entropic force with respect to the extension of the entropic spring:

$$\begin{aligned} k_{\mathrm{en},i} = \frac{\mathrm{d}F_e}{\mathrm{d}L_{\mathrm{en},i}} &= \frac{\mathrm{d}}{\mathrm{d}L_{\mathrm{en},i}} \left( \frac{k_BT}{l_P} \left( \frac{L_{\mathrm{en},i}}{l_{c,i}} + c_s + \frac{1}{4\left(1 - \frac{L_{\mathrm{en},i}}{l_{c,i}} - c_s\right)} - \frac{1}{4} \right) \right) \\ &= \frac{k_BT}{l_P} \left( \frac{1}{l_{c,i}} + \frac{l_{c,i}}{4((c_s - 1)l_{c,i} + L_{\mathrm{en},i})^2} \right) . \end{aligned} \tag{S 30}$$

The extension of the entropic spring depends on the total extension $\Delta L_{\mathrm{tot},i}$ of the filament section and the elastic spring in the filament model, assuming that the force acting on the entropic and elastic spring is the same:

$$L_{\mathrm{en},i} = \frac{k_{\mathrm{el}}}{k_{\mathrm{en},i} + k_{\mathrm{el}}} \Delta L_{\mathrm{tot},i} . \tag{S 31}$$

We rewrite the total extension of the filament section in terms of the angle $\alpha_i$ by which the

$i^{\mathrm{th}}$ filament section is deflected while it is moved down, the bead diameter $a$, the distance $\Delta y_{12}$ by which the point of interaction moves and the length $l_i$ of the $i^{\mathrm{th}}$ $(i = [1, 2])$ filament section when the filaments start to interact:

$$\Delta L_{\mathrm{tot},i} = \frac{\Delta y_{12}}{\sin \alpha_i} - \frac{a}{2} - l_i \,. \tag{S 32}$$

Substituting equation (S 31) and S 32 into equation (S 30) results in:

$$k_{\mathrm{en},i}(\Delta y_{12}) = \frac{k_B T}{l_P} \left( \frac{1}{l_{c,i}} + \frac{l_{c,i}}{4\left[(c_S - 1)l_{c,i} + \frac{k_{\mathrm{el}}}{k_{\mathrm{en},i}+k_{\mathrm{el}}}\left(\frac{\Delta y_{12}}{\sin \alpha_i} - \frac{a}{2} - l_i\right)\right]^2} \right), \; i = [1, 2] \,. \tag{S 33}$$

We solve this equation for $k_{\mathrm{en},i}$ with the Matlab function `vpasolve`. A maximum of three solutions are possible and we assume the one with solely a real part is the physically feasible solution.

In analogy to the spring constants of filament sections 1 and 2, we calculate the spring constant of filament section 3. However, the distance $\Delta y_3$ by which filament section 3 is extended is set by the the distance that beads b3 and b4 are moved and by $\Delta y_{12}$. $v$ is the speed with which the beads are moved and $t$ is the time after the interaction starts:

$$vt = \Delta y_3 + \Delta y_{12} \,. \tag{S 34}$$

Thus, the entropic spring constant of filament section 3 is:

$$k_{\mathrm{en},3}(\Delta y_{12}) = \frac{k_B T}{l_P} \left( \frac{1}{l_{c,3}} + \frac{l_{c,3}}{4\left[(c_s - 1)l_{c,3} + \frac{k_{\mathrm{el}}}{k_{\mathrm{en},3}+k_{\mathrm{el}}}(vt - \Delta y_{12})\right]^2} \right) . \tag{S 35}$$

#### 2. Deflection of the filaments

The filament sections 1 and 2 are deflected when an interaction forms and beads b3 and b4 are moved, leading to an increased contribution to the force in the $y$ direction. We calculate the projection of the forces on the $y$ axis with the factor $c_{g,i}$:

$$c_{g,i}(\Delta y_{12}) = \sin\alpha_i(\Delta y_{12}) = \sin\left(\arctan\left(\frac{\Delta y_{12}}{l_i + a/2}\right)\right) = \frac{\frac{\Delta y_{12}}{l_i+a/2}}{\sqrt{\left(\frac{\Delta y_{12}}{l_i+a/2}\right)^2 + 1}}\,, \tag{S 36}$$

see Figs. S17 and S20 for definition of the parameters. The total force acting on the interaction site in the $y$ direction therefore is:

$$F = \Big(c_{g,1}(\Delta y_{12})k_{f,1}(\Delta y_{12}) + c_{g,2}(\Delta y_{12})k_{f,2}(\Delta y_{12})\Big) \cdot \Delta y_{12}\,. \tag{S 37}$$

The forces in the $y$ direction are balanced, $F = F_3$, and by using equations (S 34) and (S 37) we thus obtain:

$$(c_{g,1}(\Delta y_{12})k_{f,1}(\Delta y_{12}) + c_{g,2}(\Delta y_{12})k_{f,2}(\Delta y_{12})) \cdot \Delta y_{12} = (vt - \Delta y_{12}) \cdot k_3(vt - \Delta y_{12})\,. \tag{S 38}$$

We solve equation (S 38) with the Matlab function `fminsearch`. A typical, calculated force increase is shown in Fig. S20c, and it qualitatively resembles the experimental data. The calculated force increase is well-described by an exponential function for small forces and a subsequent linear increase.

#### 7.3 Experimental force increase

The results of the microscopic model as well as the experimental force-time curves are well represented by a simplified description consisting of an exponential increase at low forces and a linear increase at times $t \geq t_c$. Therefore, this simplified description is used to fit the experimental data, curves that reach forces > 10 pN are fitted by:

$$F(t) = \begin{cases} f(t) = A\exp(Bt) & t \leq t_c \\ g(t) = f(t_c)\,(1 + B(t - t_c)) & t \geq t_c\,, \end{cases}$$

utilizing that $F(t_c)$ is continuous and differentiable. The parameters $A$ and $B$ are constants. Experiments where the interaction ruptures at < 10 pN are fitted by $f(t)$. We perform this analysis for all initial increases that start at zero force. For re-binding events under force (kinks), the increase is assumed to be equal to the initial increase in the same measurement.

#### 7.4 Comparison of simulated and experimental data

To determine $r_{e,u}$ and $x_u$, equations (S 24) and (S 25) are solved for different pairs of $r_{e,u}$ and $x_u$ with the force increase $F(t)$ derived from the experimental data. $r_{e,u}$ is varied from 0.01 s$^{-1}$ to 0.5 s$^{-1}$ in steps of 0.01 s$^{-1}$ and $x_u$ is varied from 0 nm to 0.8 nm in steps of 0.01 nm. The resulting force distributions for all experimental force increases and a specific $r_{e,u}$–$x_u$ pair are averaged and compared to the experimental force histogram with the Kolmogorov-Smirnov test [18]. If the simulated and experimental histograms differ less than allowed by the 5% significance level of the Kolmogorov-Smirnov test, the $r_{e,u}$–$x_u$ pair is accepted. We start with the evaluation of $r_{e,u}$-$x_u$ pairs around $r_{e,u} = 0.1$ s$^{-1}$ and $x_u = 0.1$ nm and evaluate only $r_{e,u}$–$x_u$ pairs around previously accepted $r_{e,u}$-$x_u$ pairs. We determine the values of $r_{e,u}$ and $x_u$ at the centroid of the valid parameter space, $\overline{r_{e,u}}$ and $\overline{x_u}$, shown in Fig. 3d in the main text.

#### 7.5 Calculation of the energy landscape

To calculate the energy landscapes in Fig. 3e,f in the main text, we calculate the equilibrium free energy difference $\Delta G$ from the force-independent binding, $r_{e,b}$, and unbinding rates at the centroid of the valid parameter space, $\overline{r_{e,u}}$:

$$\Delta G = -k_B T \ln \left( \frac{\overline{r_{e,u}}}{r_{e,b}} \right) . \quad \text{(S 39)}$$

The absolute values of the activation energies cannot be determined from our experiments. However, we can calculate the differences between activation energies between different conditions, *e.g.* condition $M$ ($c(\mathrm{MgCl_2})$=5 mM) and condition $T$ $c$(TX)=0.16 mM):

$$\frac{r_{e,b,M}}{r_{e,b,T}} = \frac{\exp\left(\frac{-E_{Ab,M}}{k_B T}\right)}{\exp\left(\frac{-E_{Ab,T}}{k_B T}\right)} \Rightarrow k_B T \ln \left( \frac{r_{e,b,M}}{r_{e,b,T}} \right) = E_{Ab,T} - E_{Ab,M} . \quad \text{(S 40)}$$

We set the sum $x_c$ of $x_b$ and $x_u$ to $x_c = 2$ nm as shown in Fig. 3e,f in the main text. The exact value of $x_b$ in the range of $0.4 - 2$ nm does not change the resulting force histograms, since $r_{e,b}$ dominates the force-dependent term of equation (S 24). From $\overline{x_u}$ and $x_c$, we calculate $x_b$. We directly plot $x_b$ and $x_u$ in the energy landscape as the distances from the bound or unbound state to the transition state.

#### 8 Supplementary movies

Movie S1: Movie of the interaction experiment presented Fig. 2 in the main text.

Movie S2: A strong interaction leading to zipping and the rupture of the interaction, as shown in Fig. S18.

#### References

[1] J. Block, V. Schroeder, P. Pawelzyk, N. Willenbacher, and S. Köster, "Physical Properties of Cytoplasmic Intermediate Filaments," *Biochim. Biophys. Acta, Mol.*

*Cell Res.*, vol. 1853, no. 11, pp. 3053–3064, 2015.

[10] M. Schopferer, H. Bär, B. Hochstein, S. Sharma, N. Mücke, H. Herrmann, and N. Wil-

lenbacher, "Desmin and Vimentin Intermediate Filament Networks: Their Viscoelastic Properties Investigated by Mechanical Rheometry," *J. Mol. Biol.*, vol. 388, no. 1, pp. 133–143, 2009.

[11] B. Nöding and S. Köster, "Intermediate Filaments in Small Configuration Spaces," *Phys. Rev. Lett.*, vol. 108, p. 088101, 2012.

[12] A. V. Schepers, C. Lorenz, and S. Köster, "Tuning Intermediate Filament Mechanics by Variation of pH and Ion Charges," *Nanoscale*, vol. 12, no. 28, pp. 15236–15245, 2020.

[13] G. I. Bell, "Models for the specific adhesion of cells to cells," *Science*, vol. 200, no. 4342, pp. 618–627, 1978.

[14] A. Kolomeisky, *Motor Proteins and Molecular Motors.* CRC Press, 2015.

[15] L. Schaedel, C. Lorenz, A. V. Schepers, S. Klumpp, and S. Köster, "Vimentin Intermediate Filaments Stabilize Dynamic Microtubules by Direct Interactions," 2020. bioRxiv 10.1101/2020.05.20.106179.

[16] J. Block, H. Witt, A. Candelli, E. J. G. Peterman, G. J. L. Wuite, A. Janshoff, and S. Köster, "Nonlinear Loading-Rate-Dependent Force Response of Individual Vimentin Intermediate Filaments to Applied Strain," *Phys. Rev. Lett.*, vol. 118, p. 048101, 2017.

[17] J. F. Marko and E. D. Siggia, "Stretching DNA," *Macromolecules*, vol. 28, no. 26, pp. 8759–8770, 1995.

[18] G. W. Corder and D. I. Foreman, *Nonparametric Statistics: A Step-By-Step Approach.* John Wiley & Sons, 2014.
