## Supplementary figures and images for "Multiscale mechanics and temporal evolution of vimentin intermediate filament networks"

### Supplementary movie 1

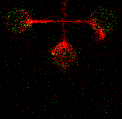

### Supplementary movie 2

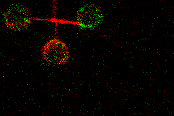
